# The differentiation of myeloid progenitors is effected by cascading waves of coordinated gene expression that remodel cellular physiology in a characteristic sequence

**DOI:** 10.1101/2025.05.25.656046

**Authors:** Andrea Repele, Joanna Handzlik, Nimasha Samarawickrama, Yen Lee Loh, Manu

## Abstract

The differentiation of hematopoietic progenitors into specialized types requires the transmittal of information from a few external or internal regulators to the thousands of genes that produce a cell type’s characteristic phenotypes. While the main signaling pathways, transcription factors, and the genes eliciting the terminal phenotypes are known, how information flows from a few regulators to thousands of genes to change the state of the cell remains to be fleshed out. To profile this information transfer process, we sampled the differentiation of the PUER myeloid cell line into macrophages and neutrophils at 29 time points over seven days. There is extensive transient regulation; the number of transcripts modulated in time is twice the number differentially expressed between endpoints. Differentiation is marked by two sharp transitions, at ∼ 8h and ∼ 80h, when transcriptomic state changes suddenly. We utilized non-negative matrix factorization to identify *behaviors*, characteristic temporal patterns of gene expression, and to classify transcripts by behavior. Only 10 distinct behaviors are sufficient to recapitulate the expression of ∼36,000 transcripts with high fidelity. Gene expression in most of the behaviors occurs in pulses of varying initiation times and durations. This implies that information transfer during differentiation occurs in cascading waves of gene expression culminating in the permanent turning on of certain genes after ∼ 80h. Each behavior is enriched in specific biological processes, so that physiological remodeling proceeds in a characteristic order—signal transduction, translation and mRNA processing, metabolism, and, ultimately, myeloid phenotypic processes. The sharp transition at 8h corresponds to the completion of transcriptional and translational remodeling and the initiation of metabolic remodeling; the one at 80h corresponds to the elicitation of myeloid phenotypes. Our analysis shows that differentiation relies upon a series of transient, rapid, and complex gene regulatory events and highlights the importance of profiling it at a high temporal resolution.

**Author summary:** The maturation of hematopoietic progenitor cells into differentiated cell types occurs over a period of about a week. This process requires the progenitors to respond to external signals by changing the expression of thousands of genes to elicit the required phenotypes. We profiled how information is transferred from a few upstream regulators to thousands of genes by measuring genome-wide gene expression at high temporal resolution during white-blood cell differentiation. We show that the information transfer occurs in cascading waves, some as short as 8 hours and others lasting for 3 days, in which thousands of genes change expression coordinately. The physiological processes remodeled in each successive wave follow a characteristic order, starting with signal transduction pathways, followed by translation and mRNA processing, then metabolism, and culminating in the production of innate immunity phenotypes. Maturation is also punctuated with two sharp transitions, when genome-wide expression changes suddenly, associated with the initiation of metabolic remodeling and the production of terminal phenotypes. Our analysis shows that a complete description of differentiation requires the characterization of transient changes and not just those observable at the endpoints.

## Introduction

Macrophages and neutrophils are developmentally closely related cell types that share a common progenitor, the granulocyte-monocyte progenitor (GMP) [1, 2]. The choice of developmental fate, macrophage or neutrophil, is understood to involve well-known hematopoietic transcription factors (TFs) that regulate each other in the manner of a bistable switch [3, 4]. This decision is thought to depend on the ratio of the expression of the TF PU.1, known to be essential for the development of all white blood cell lineages [5], and C/EBP*α*, whose expression is necessary for neutrophil development [6]. A high PU.1-C/EBP*α* ratio was shown to promote the macrophage fate, while a low ratio favored neutrophil development [7]. The actual bistable switch effecting the decision is believed to be downstream of PU.1 and C/EBP*α*. One proposal involves mutual repression between the TFs Egr1/Egr2/Nab2 and Gfi1, promoting the macrophage and neutrophil fates respectively [3]. Another bistable switch may operate between Irf8 and Gfi1 [8] either in exclusion of the Egr1/Egr2/Nab2-Gfi1 switch or in complementarity of it.

While we have some insight into the key regulators that bias the initial conditions towards one fate or the other, we know relatively little about how these initial biases are converted into the ultimate phenotypic differences observable between distinct cell types. Comparisons of hematopoietic cell types typically reveals hundreds to thousands of differentially expressed genes [9]. In *in vitro* differentiation [7, 10] and transdifferentiation [11] experiments, the initial bias could be introduced by modulating the expression of a single TF. This raises several outstanding questions about how information flows from a few upstream regulators to thousands of genes to remodel the transcriptome of a terminally differentiated cell and elicit its observable phenotypes. Are the genes that produce the observable physiological phenotypes induced early or late during differentiation? When are the determinants of the identity of the cell, such as TFs and signal transduction factors, induced and, conversely, when are the determinants of the progenitor and alternative lineages downregulated? Do any genes change expression transiently and revert in expression to their state in progenitor cells? On what time scales do these changes occur? To summarize, relatively little is known about the timing, sequence, and scale of genome-wide gene expression modulation during macrophage-neutrophil differentiation [12, 13].

In order to profile the timing and sequence of gene expression changes as they snowball from a few genes to the scale of the genome, we acquired a high resolution RNA-Seq time series dataset of *in vitro* macrophage-neutrophil differentiation of PU.1 estrogen receptor (PUER) cells, an important model of myeloid development [7, 14, 15] (Fig. 1A). PUER cells are an IL3 dependent hematopoietic progenitor cell line derived from the fetal liver of PU.1*^−/−^*mice in which PU.1 has been reintroduced after fusion to the ligand binding domain of the estrogen receptor [16]. In the absence of 4-hydroxy-tamoxifen (OHT), the PUER fusion protein is inactive so that uninduced PUER cells have a PU.1*^−/−^*phenotype and can be maintained indefinitely as bipotential progenitors by culture in medium containing IL3. OHT activates the PUER fusion protein by binding to the estrogen receptor domain, restoring the function of the PU.1 TF and relieving the block on the progenitors to induce differentiation. PUER cells can be differentiated into macrophages by OHT treatment over a period of 168 hours and into neutrophils by substituting GCSF for IL3, culturing in GCSF medium for 48 hours, and subsequently treating with OHT for 168 hours. Fate transformation in PUER cells therefore, is initiated by just two inputs, the TF PU.1 and the cytokine GCSF, making the system a good candidate for observing the information transfer process in a bifurcating cell-fate decision.

**Fig 1.**
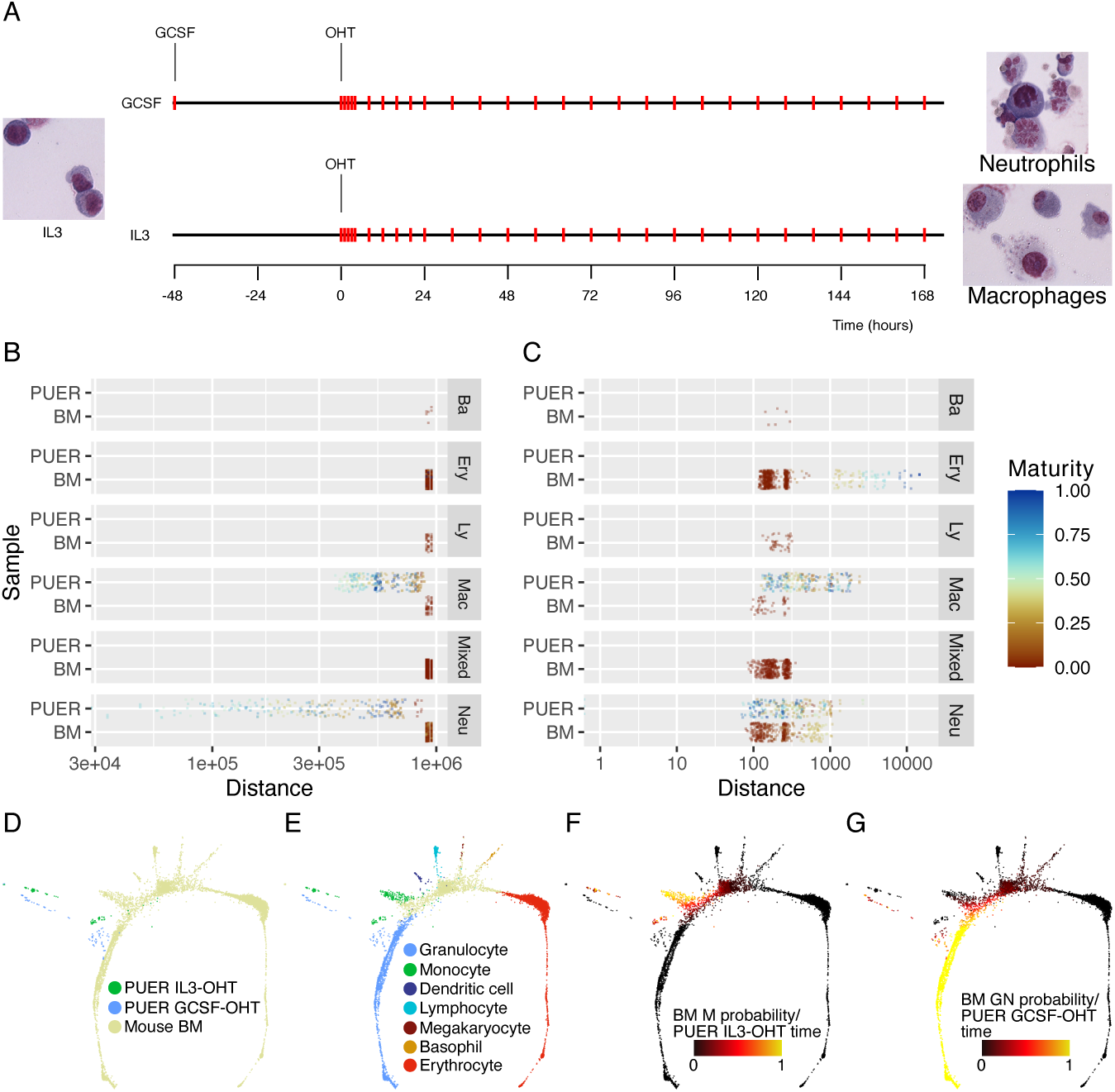
*in vitro* PUER cell differentiation reprograms PU.1*^−/−^* cells to states resembling *in vivo* bone marrow macrophage and neutrophil progenitors. **A.** Schematic showing the *in vitro* differentiation of PUER cells into macrophages and neutrophils. Red ticks indicate the timepoints at which cells were sampled for RNA-Seq. B,C The distribution of Euclidean distances between 88h GCSF-OHT samples and other PUER samples or *in vivo* BM single cells [17] either before (B) or after (C) the removal of batch effects. PUER cells in GCSF and IL3 conditions are labeled as Neu or Mac respectively. BM cells having a population balance analysis (PBA) [19] probability of more than 0.8 for a particular lineage are labeled as that lineage. Ba: basophil, Ery: erythrocyte, Ly: lymphocyte, Mac: monocyte, Neu: granulocyte. BM cells that have PBA probability less than 0.8 for all lineages are labeled as Mixed. The color of the points indicates developmental maturity so that higher more differentiated cells have higher values. For PUER cells, the developmental maturity is given by the differentiation timepoint scaled to [0, 1]. For BM cells, the developmental maturity is given by the additive inverse of the PBA potential scaled to [0, 1]. D–G SPRING [20] plots of batch-corrected PUER samples and BM cells. D. Color shows PUER samples in IL3 (macrophage) or GCSF (neutrophil) conditions and BM cells. E. Color annotates PUER samples and BM cells by lineage. Only BM cells having a PBA probability of more than 0.8 for a particular lineage are annotated. F. BM cells are annotated by a heatmap of PBA probability for the monocyte (M) lineage. IL3-OHT PUER cells are annotated by a heatmap of differentiation timepoint. G. BM cells are annotated by a heatmap of PBA probability for the granulocyte (GN) lineage. GCSF-OHT PUER cells are annotated by a heatmap of differentiation timepoint.

Our analysis of the RNA-Seq data shows that genes are expressed in diverse and complex temporal patterns throughout the course of differentiation. Differential expression analysis between the end points, while identifying thousands of genes that change in expression, underestimates the number of genes modulated in expression during differentiation since a large number of genes change expression only transiently. To overcome this limitation of pairwise comparisons, we utilized non-negative matrix factorization (NMF) to identify characteristic temporal patterns of gene expression over the entire duration of differentiation, called “behaviors”, and to cluster the genes by similarity to the behaviors. We found that only ten behaviors were sufficient to account for the temporal expression patterns of ∼36,000 transcripts and that each behavior was exhibited by hundreds to thousands of genes. Many of the behaviors were pulsatile so that expression was temporally restricted to periods ranging from 8 hours to 3 days. Gene ontology (GO) analysis showed that each behavior is enriched in specific physiological processes so that information is transferred in a characteristic order culminating in the upregulation of genes that determine terminal myeloid phenotypes at 80h. We also document two sharp transitions during differentiation, at 8h and 80h, when the global transcriptomic state changes quite suddenly. The first transition is shown to be associated with the initiation of metabolic remodeling and the second with the upregulation of myeloid phenotypic genes. Our analysis shows that information is transferred in successive cascading waves of gene expression during differentiation, punctuated by sharp transitions, each wave remodeling different aspects of the cells’ physiology in a characteristic order.

## Results

### Differentiation reprograms PUER cells toward states resembling *in vivo* bone-marrow myeloid cells

PUER cells were isolated nearly three decades ago from the bone marrow of PU.1*^−/−^* mice and resemble macrophages and neutrophils morphologically, and in the expression of select markers, when differentiated [3, 7, 15]. The bulk RNA-Seq data reported here, along with publicly available single-cell RNA-Seq (scRNA-Seq) datasets, presented an opportunity to quantitatively compare the transcriptomic state of differentiating PUER cells to those developing *in vivo*.

We chose an scRNA-Seq dataset of about 5,000 Kit+ mouse bone-marrow (BM) progenitors [17] to compare with the PUER data. The two datasets were acquired completely independently using starkly different experimental methodologies and a direct comparison is not possible. We registered the two datasets to each other by removing batch effects using MNNCorrect [18] treating each PUER sample as if it were a single cell (see Methods). Euclidean distance after cosine normalization of the genome-wide gene expression vectors [18] was used as a measure of the dissimilarity of transcriptomic state between samples or cells. Prior to batch correction, PUER samples are equidistant from all the different BM cell types even though the distances gradually increase with time within the PUER dataset (Fig. 1B). This suggests that PUER samples are very distant from BM cells, potentially due to batch effects. After batch correction, the distances between PUER samples and BM cells vary over the same range as the distances between different PUER samples (Fig. 1C), so that many PUER samples are closer to BM cells than they are to other PUER samples.

After batch correction, BM cells are among the 10 nearest neighbors of many, but not all, differentiated PUER samples (Fig. 1C). Tusi *et al.* [17] used cell-fate specific gene expression signatures to assign each cell a probability to developing a particular cell fate. 88h after OHT induction in GCSF conditions (neutrophil differentiation) PUER samples are closest to BM cells with neutrophil probability greater that 0.8. Reflecting the hierarchical cell-fate relationships during hematopoiesis, the next closest cells are BM macrophages, followed by BM basophils, BM lymphocytes, and the most distant are BM erythrocytes.

We utilized SPRING plots [21], force-directed layouts of k-nearest neighbor (KNN) graphs of cells or samples, to visualize the overall relationship of differentiating PUER cells to BM cells. In the SPRING layout, differentiating PUER samples are placed at varying distances from the BM cells (Fig. 1D). If the BM cells having more than 80% probability of achieving a particular fate are annotated, the PUER samples are closest to the macrophage/neutrophil junction, with GCSF-OHT PUER cells closest to BM neutrophils and IL3-OHT PUER cells closest to BM macrophages (Fig. 1E). When the plot is annotated with measures of cell differentiation, time for PUER samples and fate probability for BM neutrophils (Fig. 1F) or BM macrophages (Fig. 1G), the undifferentiated and early time point PUER samples are the most distant from BM cells. PUER cells progressively approach BM cells at later time points. Furthermore, PUER cells approach BM cells having a neutrophil probability close to 1 during neutrophil differentiation and BM cells having macrophage probability *>* 0.6 during macrophage differentiation.

These interrelationships between differentiating PUER cells *in vitro* and mouse BM cells developing *in vivo* support three conclusions. First, undifferentiated PUER samples are distant in their transcriptomic state from any BM cell, including progenitors. This can be understood to be a consequence of undifferentiated PUER cells being PU.1*^−/−^*mutants. Second, upon OHT induction, which rescues the PU.1*^−/−^* mutant, PUER cells approach BM cells in their transcriptomic state. Third, differentiated PUER neutrophils and macrophages are closer in their transcriptomic state to neutrophils and macrophages developing *in vivo* respectively than to any other BM cell type. The differentiation of PUER cells can thus be seen as the reprogramming of cells from a mutant state to one very close to wildtype BM myeloid cells.

### Global changes in the gene expression landscape during myeloid differentiation

As a first step we sought to uncover temporal patterns of changes in genome-wide gene expression. We computed Pearson correlation of genome-wide gene expression between all pairs of time points (Fig. 2A,B). As one would expect, the correlation coefficient is higher for nearby time points and reduces as the difference between the time points increases. The largest effect is that of GCSF pre-treatment (−48h vs. 0h; Fig. 2A), which can be understood as the result of the large time difference of 48 hours and the important role that GCSF plays in the growth and maturation of hematopoietic cells [22]. Unexpectedly, we found that time points could be divided into at least two groups, early (*<* 12 hours) and late (≥ 12 hours), so that timepoints have much higher correlation within each group than to timepoints from the other group. This is most easily discerned in the macrophage differentiation in IL3 conditions (Fig. 2B). For example, the 16h timepoint has very low correlation, 0 ≥ *r <* 0.2, with the 2h timepoint in the early group, occurring only 14 hours earlier, but has very high correlation, *r >* 0.8, with the similarly spaced 32h timepoint in the late group. Although post-OHT timepoints have higher correlation overall, a similar pattern is discernible in the GCSF condition. That the differentiation can be divided into two phases suggests that there is a large-scale transition in genome-wide gene expression patterns occurring around 8 − 12 hours.

**Fig 2.**
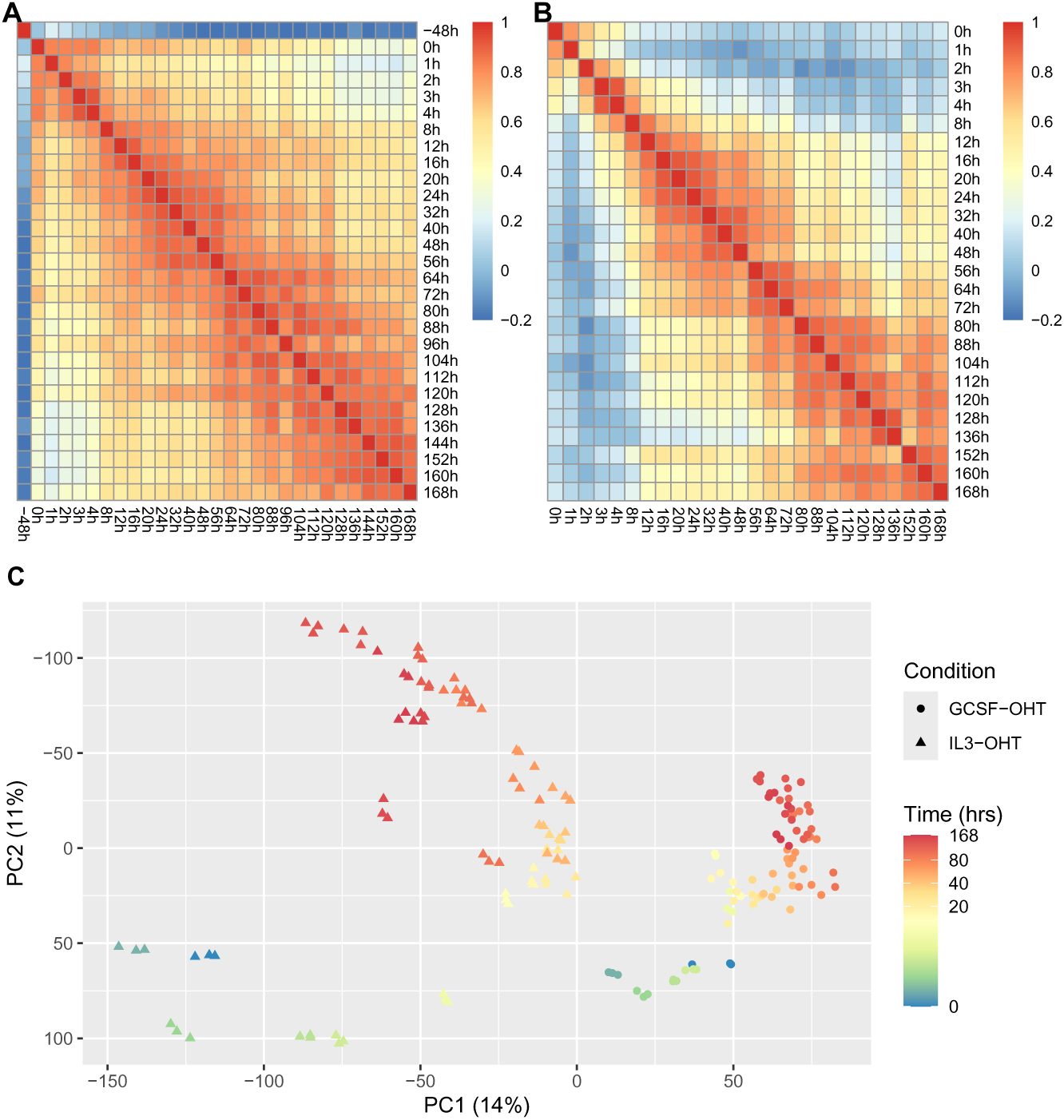
PUER cell differentiation is punctuated by two sharp transitions. **A,B**. Pearson correlation coefficient between genome-wide gene expression at each pair of time points in GCSF (**A**) or IL3 (**B**) conditions is plotted as a heatmap. **C**. Principal components analysis of all the samples. Gene expression was standardized to have zero mean and unit variance. The samples are plotted along the first two principal components, PC1 and PC2, that account for 25% of the total variance.

We further characterized the global patterns of gene expression time evolution using principal component analysis (PCA) (Fig. 2C). The first two principal component axes accounted for 45% of the total variance in the data, hinting that there are two main effects and that the two-dimensional space is a good approximation to the high-dimensional gene expression space. The GCSF and IL3 conditions follow distinct differentiation trajectories and are separated by a large shift along the PC1 axis, occurring during the GCSF pre-treatment, which implies that the first principal component corresponds to the effect of GCSF. In both the GCSF and IL3 conditions, the trajectories move along the PC2 axis after OHT addition at 0h, suggesting that the second principal component corresponds to the effect of PU.1 and time. The displacement between the 8h and 12h IL3 time points is the second largest after that of GCSF pre-treatment, which implies a very large rate of change in genome-wide gene expression given that it occurs in only 4 hours compared to the 48 hour duration of GCSF pre-treatment. Similar but smaller jumps are observed between the 4h and 8h GCSF and the 72h and 80h IL3 time points, while there is a sharp reversal of the direction of movement at the 136h IL3 timepoint. These jumps corroborate the inference drawn from the Pearson correlation analysis (Fig. 2A,B) that there is a large-scale transition in the pattern of genome-wide gene expression around 8 – 12 hours after OHT induction, and indicate that there are other such transitions occurring at later stages of the differentiation as well.

### Diversity of temporal patterns of transient gene expression

In order to gain insight into the genome-wide transitions occurring during the course of differentiation we next analyzed the temporal patterns of the expression of individual genes. We enriched for genes likely to be regulated during the differentiation process by first identifying genes expressed differentially between the end points, undifferentiated PUER cells, −48h GCSF or 0h IL3, and 7-day OHT treated samples, 168h GCSF or IL3 (Fig. 3A,D). 43% and 27% of genes were differentially expressed between the end points in GCSF and IL3 conditions respectively. The neutrophil differentiation is the compounded effect of GCSF and OHT treatments and one may discern between the two by identifying genes differentially expressed because of GCSF pre-treatment, by comparing −48h samples to 0h GCSF samples, and those differentially expressed before and after 7 days of OHT treatment, by comparing 0h to 168h GCSF samples (Fig. 3B,C). Consistent with both the correlation analysis and PCA, more genes are differentially expressed due to GCSF pre-treatment (37%) than due to OHT treatment (29%), even though the latter is conducted over a larger time interval. There is significant overlap in the genes differentially expressed in the two conditions with common DEGs comprising 73% and 44% of all the genes differentially expressed between IL3 endpoints and GCSF endpoints respectively (Fig. 3E,F).

**Fig 3.**
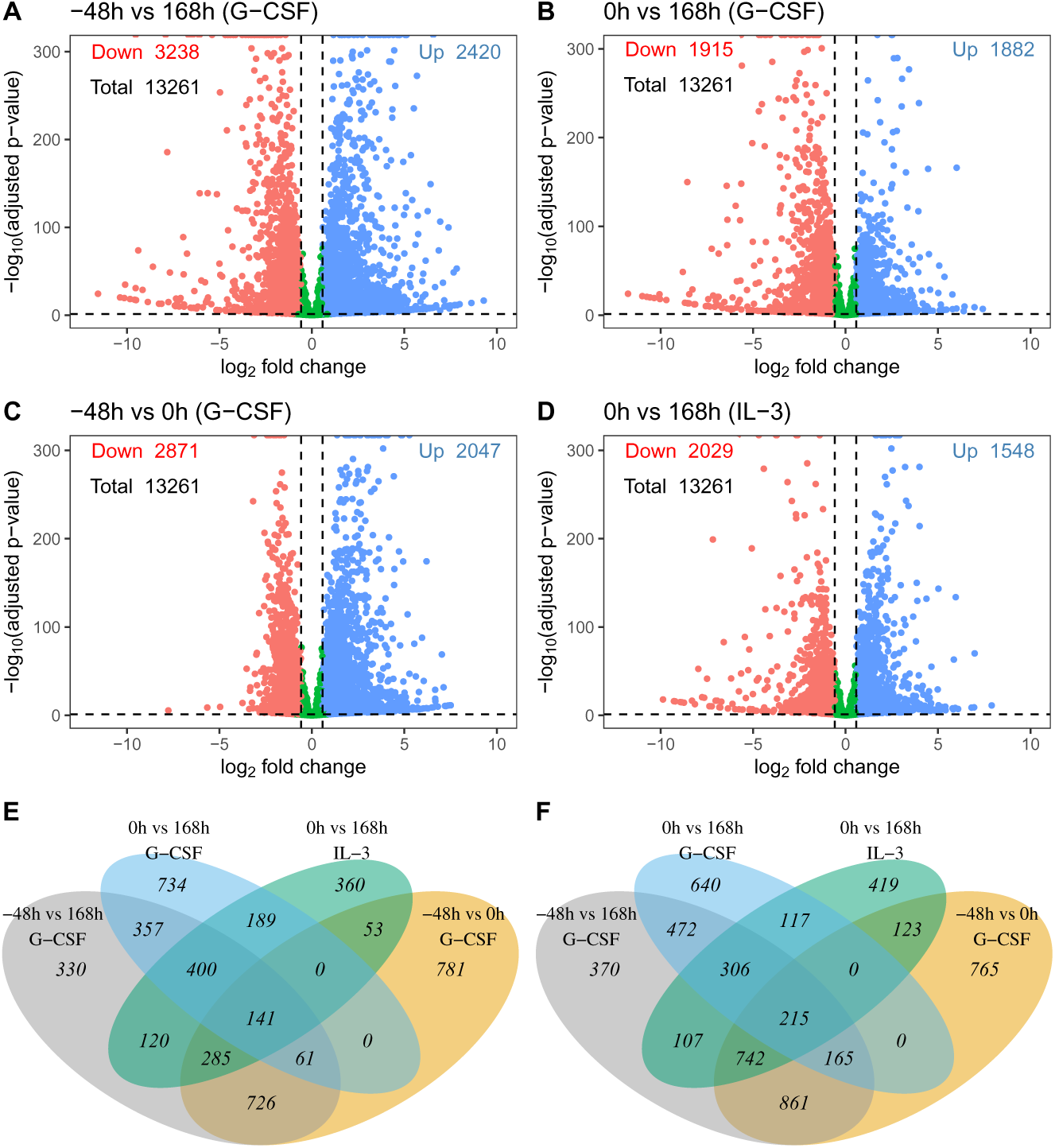
The identification of genes expressed differentially between the endpoints of the differentiation. **A–D**. Scatter plots of *p*-value vs. fold change for all the genes. The *p*-value and fold change thresholds used to identify DEGs (see Methods) are shown as horizontal and vertical dashed lines respectively. **E–F**. Venn diagrams showing the intersection of the different sets of DEGs identified in this analysis. **A**. Comparison of undifferentiated PUER cells (−48h) with cells treated with OHT for 7 days in GCSF conditions (168h). **B**. Comparison of PUER cells pre-treated with GCSF for 48 hours (0h) with cells treated with OHT for 7 days in GCSF conditions (168h). **C**. Comparison of undifferentiated PUER cells (−48h) with those pre-treated with GCSF for 48 hours (0h). **D**. Comparison of undifferentiated PUER cells (−48h) with those treated with OHT for 7 days in IL3 conditions (168h). **E,F**. Overlap of DEGs for selected time points for up-regulated (E) and down-regulated (F) genes.

The temporal expression patterns of the differentially expressed genes are very diverse and show extensive transient regulation, in which expression at the start and end of differentiation is similar but is modulated in the middle (Fig. S6). In order to better reveal broader patterns, the genes were clustered hierarchically according to the similarity of their temporal expression patterns. Several patterns are noticeable. Consistent with all the previous analyses, GCSF pre-treatment exerts significant effect on the gene expression, with a large number of genes turning off and a smaller but still sizable group turning on at 0h in the GCSF treatment (Fig. S6A). Another significant shift in the gene expression occurs around the 8 − 12h timepoints, when a large number of genes are upregulated and a smaller number of genes are downregulated. Furthermore, the number of genes coordinately regulated in this manner is greater in the IL3 condition than the GCSF condition. Also discernible in the IL3 condition, but less so in the GCSF condition, are several waves of transient gene upregulation and downregulation during the first 8 hours of differentiation, with different groups of genes peaking at different timepoints. Finally, a transition at 80h in the IL3 condition when a smaller number of genes are either up- or down-regulated can also be seen. To summarize, the temporal gene expression patterns further corroborate the transitions inferred previously, show that the first 8 − 12 hours of differentiation involve rapid changes in expression, and reveal extensive transient and coordinated regulation of groups of genes.

The above analysis ignores genes that are transiently expressed during differentiation but revert to their original expression by the end and are not identifiable as DEGs between the end points. Focusing only on DEGs between endpoints, therefore, could potentially ignore important aspects of the information transfer process of differentiation. We sought a method to identify and classify genes based on the entire temporal pattern of expression instead of the behavior at the end points. Non-negative matrix factorization (NMF) [23, 24] is an unsupervised learning method that simultaneously performs dimensionality reduction and clustering. Given the expression of gene *g* at time *t*, *X_gt_*, we used NMF to find *k* gene expression patterns or *behaviors*, *H_mt_*, and weights, *W_gm_*, so that the expression of the gene is approximated as a weighted sum of the *k* behaviors, 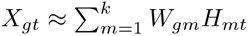. The behaviors and weights were restricted to be non-negative so that the a behavior is interpretable as the expression of a metagene. Dimensionality reduction was achieved by choosing the number of behaviors, *k*, to be much smaller than the total number of genes. One important capability of NMF is that it performs automatic feature identification; the behaviors are spatially or temporally restricted and correspond to parts of the overall pattern. NMF of photographs of images results in behaviors that are facial features such as noses and eyes. NMF on temporal gene expression patterns therefore is expected to identify temporally restricted features such as pulses and oscillations.

We found that the error between the original dataset and the NMF approximation has a local minimum at *k* = 7 (Fig. S4) and chose *k* = 10, with a slightly higher error, to ensure that the full diversity of temporal expression patterns was accounted for. NMF was performed on the entire dataset, including GCSF and IL3 differentiation so that the behaviors (Fig. 4 and Table 1) captured the constraints of a bipotential differentiation. With the exception of three behaviors, *k* = 3, 5, and 8, all behaviors showed mutually exclusive expression patterns so that behaviors regulated in the GCSF condition did were not regulated in IL3 and vice versa. In behavior 5, expression is downregulated in both GCSF and IL3 conditions over the first two time points after the initiation of treatment, which occurs 48 hours after substituting GCSF for IL3 in neutrophil differentiation and in the first two hours following OHT addition in macrophage differentiation. Behavior 8 is a pulse of expression starting around 8h after OHT addition and terminating around 80h in both conditions. Behavior 3 exhibits a pulse of expression from 2h to 48h followed by oscillatory expression lasting to the end of the differentiation.

**Fig 4.**
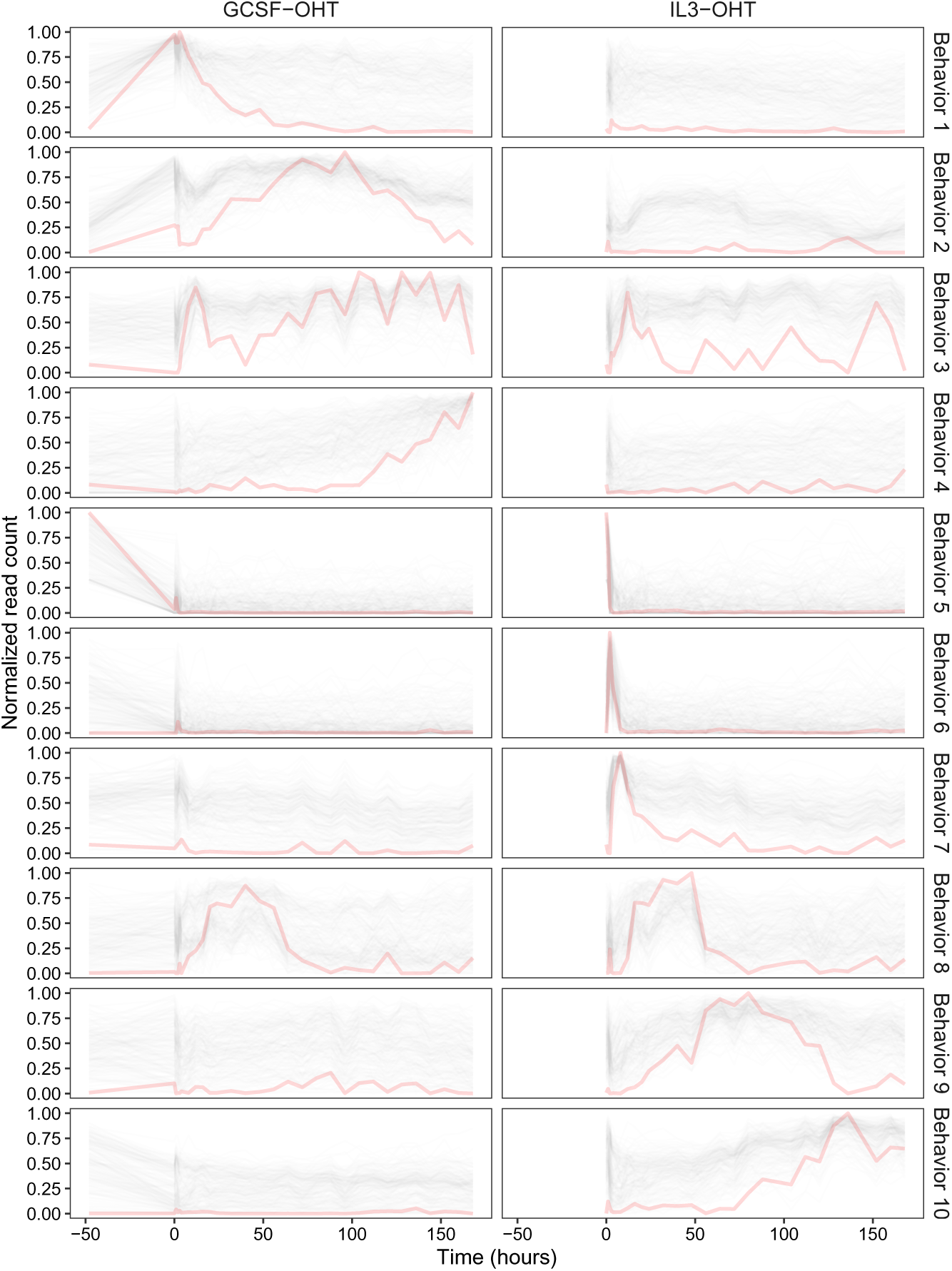
The temporal behaviors of the 10 metagenes identified by NMF. The temporal behavior of each metagene is shown in red. The temporal expression patterns, scaled between 0 and 1, of the top 200 genes for each behavior are shown in black.

**Table 1.** Summary of biological processes enriched in the top 500 genes for each behavior. The top 30 enriched terms (Fig. S11–S20) are summarized. Terms in italics were not in the top 30.

| Metagene | Behavior in GCSF | Behavior in IL3 | Enriched biological processes |
| --- | --- | --- | --- |
| 1 | Upregulated by GCSF and downregulated by OHT | Not regulated | Cell cycle, metabolism |
| 2 | Upregulated by GCSF, transiently downregulated by OHT, peaking around 100h | Not regulated | Cell cycle |
| 3 | Pulse from 2h to 48h, re-upregulated and expressed in an oscillating pattern until the end | Pulse from 2h to 48h, oscillations at later timepoints | Autophagy, metabolism, cytoskeleton, NF-kB signaling |
| 4 | Upregulated after 100h and peaks at the end | Not regulated | Granulocyte/neutrophil migration/chemotaxis, innate immune response |
| 5 | Downregulated by GCSF pre-treatment | Downregulated by OHT by 2h | Extracellular signaling, epithelial-to-mesenchymal transition, thermogenesis, cell adhesion, cytoskeleton, migration |
| 6 | Not regulated | Pulse from 0h to 8h | TGFb, Erk1/2, Ras signaling, cytoskeleton, extracellular matrix |
| 7 | Not regulated | Pulse from 2h to 48h | Translation, mRNA processing |
| 8 | Pulse from 8h to 80h | Pulse from 8h to 80h | Metabolism, leukocyte activation |
| 9 | Not regulated | Pulse from 12h to 128h | Metabolism |
| 10 | Not regulated | Upregulated after 80h and peaks at the end | Translation, metabolism, *myeloid/leukocyte proliferation, innate immunity* |

Three behaviors, *k* = 1, 2, and, 4, have expression during neutrophil differentiation but not during macrophage differentiation. Behavior 1 expression is upregulated in response to GCSF pre-treatment but is then downregulated upon OHT induction. Behavior 2 is also upregulated by GCSF and transiently downregulated by OHT, but is re-upregulated once more, peaking at 100h. In behavior 4, expression initiates around 100h and remains on until the end of the differentiation. Three macrophage specific behaviors, *k* = 6, 7, and, 9, have pulsatile expression lasting between 0 − 8h, 2 − 48h, and 12 − 128h respectively. Behavior 10 has expression that initiates around 80h and remains on until the end of the differentiation. The process of information transfer during differentiation thus appears to involve multiple successive waves of gene expression with handover occurring at time points that correspond approximately to the global transitions at 8 − 12h (behaviors 6, 8, 9) and 80 − 100h (behaviors 4, 9, 10) and culminating in the switching on of a subset of genes permanently.

It is possible that these behaviors do not correspond to parts of the expression of any one gene but are used by NMF to construct entirely different patterns by linear combination. We checked whether this was true by visualizing genome-wide gene expression after sorting transcripts according the weights of their dominant behavior (Fig. 5). About 8,000 transcripts are not expressed at all and have zero weight for all the behaviors. Another group of transcripts are expressed but their expression does not vary over time. These transcripts—particularly enriched among behavior 2 and 3 clusters—have high weights for multiple behaviors. Excepting the unexpressed and constantly expressed groups, the remaining ∼20,000 transcripts are dominated by a few behaviors so that their expression patterns match one or more behaviors. The behaviors inferred by NMF, therefore, are exhibited as real observable gene expression patterns.

**Fig 5.**
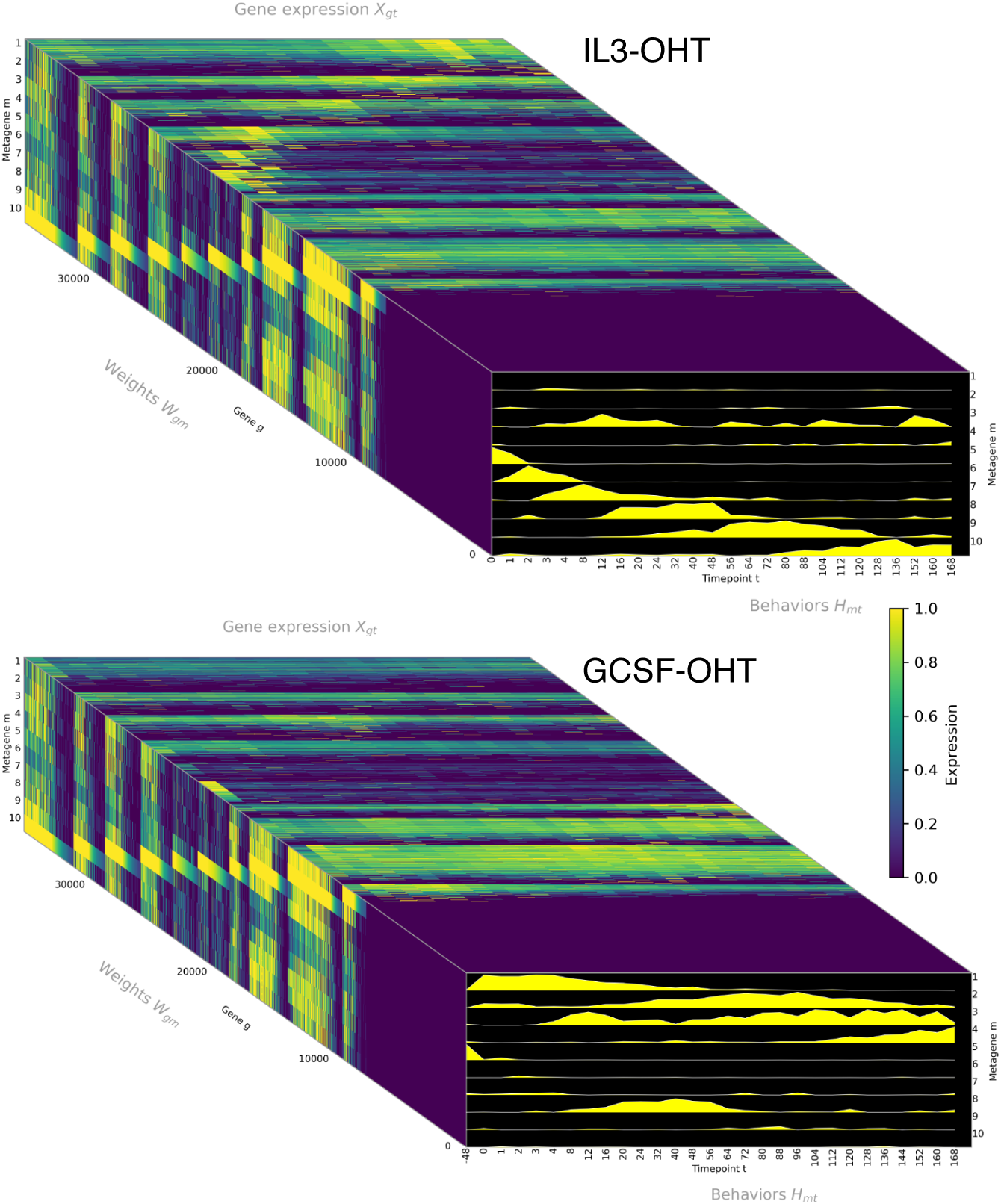
Clustering of genome-wide gene expression by temporal behaviors. The gene expression matrix (*X_gt_*), the temporal behaviors of the 10 metagenes (*H_mt_*)), and the weights of each transcript for each behavior (*W_gm_*) are depicted on the faces of a rectangular prism. *X_gt_* and *W_gm_* are heatmaps with color representing scaled gene expression or weights respectively. The behaviors are plotted as stacked lineplots with time on the *x*-axis and expression on the *y*-axis. Transcripts having the same dominant behavior were grouped together and then sorted in descending order of the weight of the dominant behavior. IL3-OHT and GCSF-OHT samples are shown on the top and bottom prisms respectively.

### Gene expression changes most rapidly during the earliest stages of the differentiation

Temporal gene expression patterns of transcripts differentially expressed between the endpoints and the pulsatile behaviors found by NMF suggested that most of the changes in genome-wide gene expression are concentrated in the first 12 hours of the differentiation. We checked whether this is true by quantifying the pace of gene expression changes during differentiation. We determined the genes differentially expressed between consecutive timepoints and utilized the number of such genes as a measure of the rate of change (Fig. 6A). The timing of sharp gene expression shifts during the differentiation may indicate important moments when crucial lineage decisions are made. Most DEGs between consecutive time points are detected before the 12h timepoint in both conditions. This confirms that rapid, early changes during the first 12 hours after OHT induction are a general feature and not just restricted to genes differentially expressed between the endpoints of the differentiation. Additionally, there is very little overlap between DEGs detected between different pair of consecutive timepoints, implying that genes are undergoing rapid but short-lived, transient changes during the early stages of the differentiation (Fig. 6B,C).

**Fig 6.**
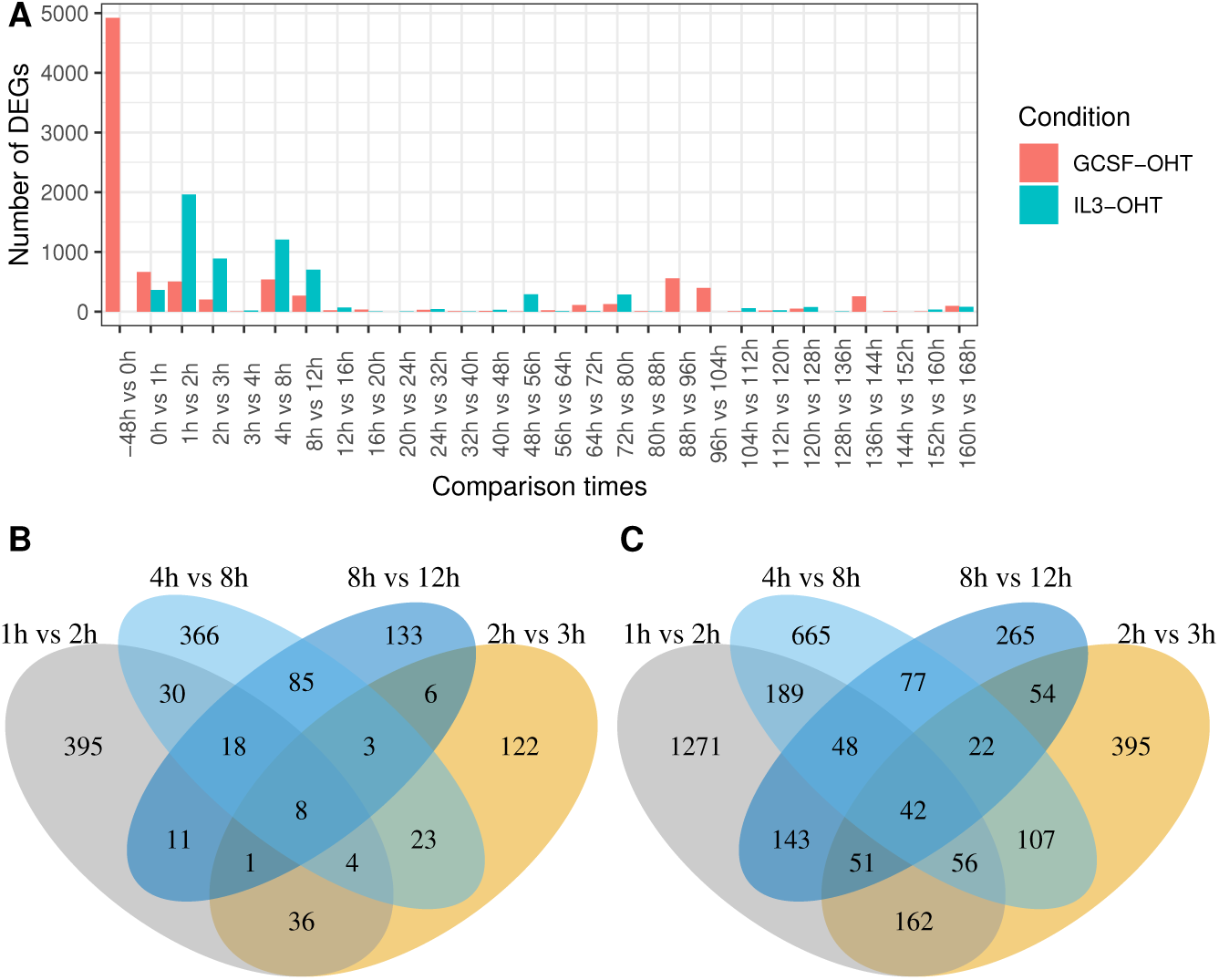
The number of DEGs between consecutive time points. (A) The number of DEGs detected between each pair of consecutive time points. The overlap between the DEGs detected at different time points in the (B) GCSF or the (C) IL3 conditions.

### Functional enrichment analysis

We performed functional enrichment analysis to determine the biological processes induced during differentiation. In one approach, we performed this analysis for DEGs detected between consecutive early time points. In a complementary approach, we determined gene ontology (GO) terms enriched for the top 500 genes for each of the ten behaviors.

#### DEGs between consecutive early time points

The GO terms enriched in the DEGs ascertained between consecutive early time points (Figs. 7, 8, S7–S10) show clear temporal patterns. The top 30 GO terms—having the lowest 30 adjusted *p*-values—enriched in the first comparison, 0h vs. 1h (Fig. 7 and 8), are dominated by terms for intracellular signaling relating to MAPK, Erk1/2, and other pathways in both IL3 and GCSF conditions. This suggests that PU.1 activation by OHT very rapidly induces signaling cascades. In the 1h vs. 2h comparison, there are fewer signaling-related terms while terms related to development and cellular differentiation, especially leukocyte differentiation, are prominent. This suggests that the cell-fate decision is made very early during the differentiation process. Although biological functions related to myeloid differentiation are common, GO terms related to the differentiation of non-myeloid cell lineages are also enriched, reflecting the pleiotropic roles that most hematopoietic TFs play in multiple lineages.

**Fig 7.**
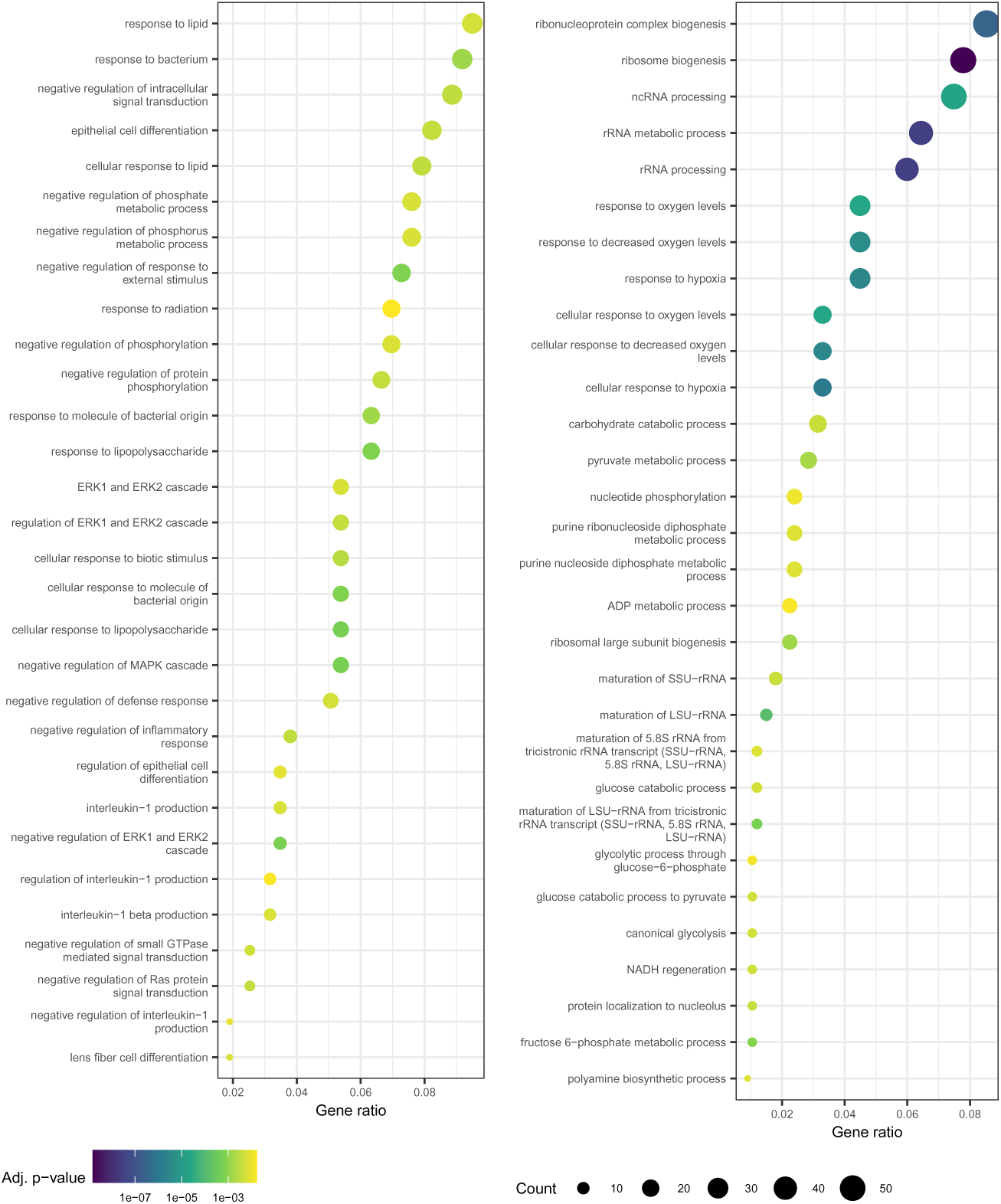
GO terms enriched for genes differentially expressed between. 0**h and** 1**h and** 8**h and** 12**h IL3 samples.** GO terms enriched for DEGs between 0h and 1h and between 8h and 12h are shown on the left and right respectively. Benjamini-Hochberg adjusted *p*-value each enriched GO term is indicated by the color of the point. The number of DEGs overlapping with a term are depicted with the size of the point. The *x*-axis is the fraction of the DEGs overlapping with a GO term (gene ratio).

**Fig 8.**
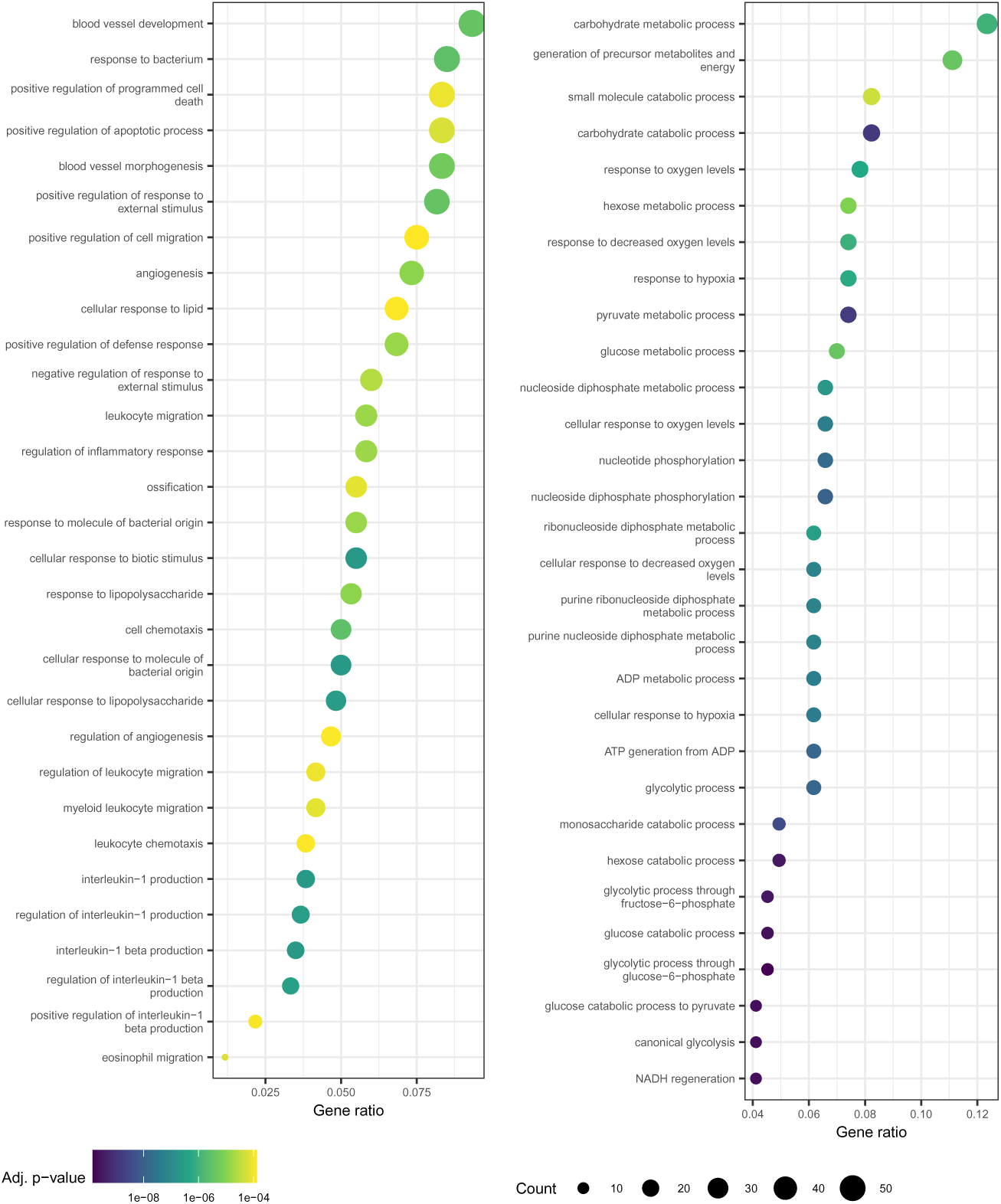
GO terms enriched for genes differentially expressed between. 0**h and** 1**h and** 8**h and** 12**h GCSF samples.** GO terms enriched for DEGs between 0h and 1h and between 8h and 12h are shown on the left and right respectively. See the legend of Fig. 7 for plot description.

The GO terms related to signal transduction, development, and cellular differentiation are no longer present in the top 30 for comparisons between 4h and 8h (Fig. S9 and S10) or between 8 and 12h (Fig. 7 and 8). Instead, GO terms related to physiological processes such metabolism, ribosome biogenesis, ion homeostasis, and cell-cell adhesion were highly ranked. This suggests that the cells initiate the remodeling of physiological processes as early as 8h in the course of the differentiation to implement a decision that’s already been made. Furthermore, this analysis identifies the 8 − 12h transition observed above (Figs. 2 and S6) as the initiation of large-scale physiological changes in response to differentiation cues. Finally, comparisons between later timepoints were also not enriched in myeloid differentiation and cell-fate commitment GO terms but were instead linked to various physiological processes, strengthening the interpretation that the cell-fate decision is made within the first 4 hours of OHT induction.

#### Top 500 genes for each behavior

The comparison of consecutive time points is limited to the detection of fast changing biological processes since genes changing expression at a slower pace may only build up statistically significant differences over larger time spans. In order to understand the progression of biological phenomena over the entire span of the differentiation, we performed GO analysis on the genes having highest 500 weights for each behavior (Fig. S11–S20, Table 1).

Mirroring the inference of the DEG GO analysis, the earliest acting behavior in IL3, behavior 6, expressed in a pulsatile manner during the first few hours, is enriched in signal transduction GO terms. The next wave of gene expression lasts from 2h to 48h (behavior 7) and involves translation and mRNA processing. This is followed by a third wave lasting from about 8h to 128h (behaviors 8 and 9) that is enriched in GO terms for metabolism and leukocyte activation. The process culminates in a set of genes that are upregulated around 80h and peak in expression at the end of the differentiation. This final IL3 wave is enriched in GO terms for myeloid proliferation and innate immunity. The GCSF condition behaviors follow a pattern similar to that of IL3. Behavior 3, which features a wave of expression from 2h to 48h followed by re-upregulation and expression until the end of the differentiation, is enriched in extracellular signaling GO terms, similar to the early pulse behavior 6, and also metabolism terms, similar to late behaviors 9 and 10 in IL3. Behavior 8, a wave of expression from 8h to 80h, is enriched in metabolism and leukocyte activation terms. A final wave of gene expression starting at about 100h and peaking at the end of the differentiation is enriched in granulocyte phenotypes such as migration, chemotaxis, and immune response.

Besides these successive waves of gene expression induced by OHT treatment, behavior 5 features genes that are downregulated in response to both GCSF and OHT treatment. This behavior is enriched in an eclectic set of terms, extracellular signaling, epithelial-to-mesenchymal transition, thermogenesis, cell adhesion, cytoskeleton, and migration. This might reflect the suppression of non-myeloid cell fates by GCSF and PU.1. Two behaviors specific to GCSF, 1 and 2, which features genes that are transiently upregulated in response to GCSF and downregulated by OHT are highly enriched for cell cycle GO terms, implying that the two signals have opposing effects on the cell cycle.

## Discussion

Time series datasets from differentiation experiments [10] are a powerful method for inferring GRNs and clarifying the causality of gene regulatory events during development [25]. While such data are acquired *in vitro* and may be only an approximation to the *in vivo* phenomena, they provide a window into dynamics that is not available in static snapshots of developmental processes. While pseudotime approaches [17, 21] have been used to infer the developmental sequence of cells in scRNA-Seq data, it is impossible to determine the rate of change of expression. Furthermore, *in vitro* differentiation allows one to conduct carefully controlled experiments to minimize the effect of animal-to-animal and technical variation. Despite these strengths, high-resolution time series datasets are fairly uncommon and we are only aware of one other dataset [10] of comparable temporal resolution that has been published so far in hematopoiesis.

Our analyses suggest that there is a large-scale sudden change in gene expression, reminiscent of phase transitions, occurring around 8 − 12h after the induction of PU.1 by OHT. We confirmed this conclusion in multiple different analyses: the Pearson correlation of genome-wide gene expression (Fig. 2A,B), PCA (Fig. 2C), and expression patterns of differentially expressed genes (Fig. S6). Furthermore there is another transition occurring around 80h. Similar to our data, it has been observed that there is relatively low rate of change between 12h and 48 hours and from 72h to 168h in the reprogramming of B cells into macrophages by the enforced expression of *Cebpa* [26]. This suggests perhaps that these transitions are a general phenomenon of reprogramming and not an idiosyncrasy of the PUER system. It wasn’t clear whether the jumps between 0h and 12h and 48h and 72h during B cell transdifferentiation were the result of the relatively large time intervals, 12h and 24h, or a significantly higher velocity of gene expression change. The much higher temporal resolution of our data unambiguously establishes that the jumps are the result of increased velocity—the shift between 8h and 12h in IL3 conditions is the second largest shift after the one induced by GCSF pre-treatment but occurs in 4h instead of 48h (Fig. 2C).

The trajectories of the DEGs between the endpoints of differentiation suggested extensive transient regulation so that the change in expression from undifferentiated cells peaks during the course of differentiation rather than at the end (Fig. S6). We also observed that each gene was not expressed in a unique temporal pattern; instead there were a relatively small number of patterns in each of which thousands of genes were expressed. These observations motivated two questions. First, are there genes that are not detectable as DEGs between the endpoints of differentiation but are nevertheless transiently regulated? An affirmative answer to this question would expand the set of genes participating in the information transfer process during differentiation. Second, what number and type of temporal expression patterns are sufficient to describe the changes in gene expression genome wide? Dimensionality reduction would make it easier to appreciate the larger scale patterns during differentiation.

NMF revealed that only 7–10 patterns or behaviors were sufficient to recapitulate genome-wide expression with high fidelity. To a large extent, individual transcripts exhibited these behaviors (Fig. 5) so that the behaviors reflect actual regulatory events during differentiation. Confirming our suspicions, approximately 15,000 transcripts have transient expression whereas only about 8,000 are differentially expressed between endpoints (Fig. 3). The transient expression patterns are of a specific type; OHT treatment causes thousands of genes to be expressed in a pulsatile fashion with varying pulse initiation times and durations (Table 1). This implies that information transfer during differentiation occurs in coordinated waves of gene expression culminating in the permanent turning on of certain genes after ∼ 80h. Furthermore, the time span of the pulses matches the sharp transitions observed in the transcriptome-wide data (Fig. 2). The 8 − 12h transition coincides with the termination of the behavior 6 pulse and the initiation of the behavior 8 and 9 pulses. The transition at 80h corresponds to the termination of behavior 8 and 9 pulses and the initiation of behavior 4 and 10 that remain upregulated until the end.

GO analysis of the genes with high weights for each behavior showed that most of the behaviors specialized in specific physiological processes (Table 1 and Fig.S11–S20). This implies that physiological remodeling proceeds in a characteristic order during differentiation. The earliest event is the transient upregulation of signal transduction pathways (behavior 3 and 6) during the first 8 hours followed by translation and mRNA processing (behavior 7) from 2h to 48h, and metabolism (behaviors 8 and 9) from 8h to 128h. The last set of genes to be upregulated around 80 − 100h are enriched for myeloid phenotypic processes such as neutrophil chemotaxis and innate immune response (behaviors 4 and 10). The GO analysis of the behaviors is corroborated by the pairwise comparisons between early timepoints (Fig. 7, 8, S7–S10). GO terms related to signal transduction and leukocyte differentiation are enriched among the differentially expressed genes in the comparisons between consecutive time points up to 2h. However, in the comparisons between 4h and 8h and 8h and 12h, signal transduction or myeloid differentiation terms are no longer in the top 30. Taken together these observations imply that the first transition at 8h corresponds to the completion of the remodeling of signal transduction, differentiation, transcription, and translation and the initiation of metabolic rewiring. The second transition around 80h is the completion or slowing down of metabolic remodeling and the upregulation of genes producing the phenotypic properties of the cell type. A corollary of this conclusion is that the cell-fate decision appears to have been made by the time of the 8 − 12h transition; subsequent modulation of gene expression revolves around phenotypic remodeling.

This dataset is a rich resource and the analysis reported here scratches the surface of the biological insights harbored within. One of the main goals of future work would be clarifying the causality of events at a finer granularity both in time and at the level of genes. A potential way of accomplishing that would be to conduct “multi-omic” analyses by combining these data with epigenetic assays such as ATAC-Seq [27, 28]. We [28] and others [27] have demonstrated that TF binding sites are detectable at single base pair resolution in deeply sequenced ATAC-Seq data. Therefore, it is possible in principle to determine the TFs controlling each behavior by inspecting the promoters and enhancers of member genes for TF footprints. This could result in a “blow-by-blow” description of the causality of the information transfer process during cellular differentiation. Finally, the NMF analysis, besides illuminating the information transfer process, has remarkably demonstrated that 36,255 transcripts effectively behave as 7–10 metagenes. This dimensionality reduction could enable future predictive models of macrophage-neutrophil differentiation or of multiple lineages, if combined with other datasets [10, 17].

## Methods

### PUER cell culture

PUER cells were cultured according to standard procedures [29]. PUER cells were routinely maintained in complete Iscove’s Modified Dulbecco’s Glutamax medium (IMDM; Gibco, 12440061) supplemented with 10% FBS, 50*µ*M *β*-mercaptoethanol and 5ng/mL IL3 (Peprotech, 213-13).

### PUER differentiation

PUER cells were expanded in T-75 flasks prior to the initiation of differentiation. GCSF pre-treatment was initiated by washing the cells 3 times with PBS and then seeding in 48-well plates at a concentration of 5 × 10^5^ cells/ml in PUER cell culture medium in which IL3 had been replaced by 10ng/mL Granulocyte Colony Stimulating Factor (GCSF; Peprotech, 300-23). At the same time, the PUER cells destined for the macrophage differentiation were seeded in 48-well plates at a concentration of 5 × 10^5^ cells/ml in IL3 PUER cell culture medium. After 48 hours of GCSF pre-treatment, the neutrophil differentiation was commenced by the addition of 100nM 4-hydroxy-tamoxifen (OHT; Sigma, H7904-5MG). The macrophage differentiation was initiated at the same time by the addition of 200nM OHT. In this chapter, we regard time zero (0h) of differentiation as occurring just before the addition of OHT. With this starting point, the initiation of GCSF pre-treatment occurs immediately after −48h while cells pre-treated with GCSF for 48h but not yet induced by OHT are at 0h. Since both treatments start with uninduced PUER cells, the data for the −48h time point of the neutrophil differentiation and 0h time point of the macrophage differentiation are derived from the same samples and are identical. OHT rapidly converts from the Z isomer to the E isomer, having 100-fold lower activity, in cell culture media. Half the medium was replaced and fresh OHT was added at 40h, 88h, and 136h in order to maintain differentiation pressure.

### Sample collection

In addition to undifferentiated PUER cells, which correspond to the −48h neutrophil and 0h macrophage timepoints, samples were collected after 48h of GCSF pre-treatment but before OHT induction (0h neutrophil). After OHT induction, in both GCSF and IL3 treatments, samples were collected every hour for the first four hours, every four hours for the first day, and every eight hours until the end of the seventh day (Fig. 1A). 4 biological replicates were collected for each timepoint. Cells for each time point were seeded into a dedicated 48-well plate so that they could be harvested without disturbing the samples for the other time points. Since the differentiation produces adherent cells, the cells were detached with trypsin using standard protocols for all timepoints after 24h. The cells were transferred into a 96-well plate, which was centrifuged at 1500rpm for 5 min, the majority of the medium was aspirated, the cell pellet was snap-frozen in liquid nitrogen, and stored at −80C until RNA extraction.

### Total RNA extraction, quality control, and spike in of ERCC standards

Total RNA was extracted on a Bio-Mek FX^P^ liquid handling workstation (Beckman Coulter) using the RNAdvance Tissue total RNA isolation kit (Beckman Coulter, A32649) in a 96-well format following the manufacturer’s protocol. Genomic DNA contamination was assessed by reverse transcribing the RNA with and without reverse-transcriptase and detecting GAPDH using qPCR. The number of additional cycles required to reach a threshold (Δ*C_t_*) was utilized to assess the fraction of genomic DNA (2*^−^*^Δ^*^Ct^*) and only samples with less than 1% genomic DNA were utilized for library preparation. The quality of the RNA was assessed by capillary gel electrophoresis on the Agilent 2100 Bioanalyzer using the Eukaryote Total RNA Nano kit (Agilent, 5067-1511). Only samples with RNA integrity numbers (RIN) greater than or equal to 9.5 were utilized, although there was only one sample with a RIN of 9.5 and the median RIN was 9.9. The concentration of RNA was determined using the Qubit fluorometer and the Qubit RNA High Sensitivity kit (Invitrogen, Q32855). With the exception of 8 samples with lower RNA yield, the samples were standardized to a mass of 1, 875*ng* in a 25*µl* volume. The samples with lower yield were standardized to a mass of 1, 437.5*ng* in a 25*µl* volume. 3.75*µl* or 2.88*µl* of a 1 : 100 dilution of the External RNA Control Consortium (ERCC) ExFold RNA Spike-In mix (Invitrogen, 4456739) was added to the high- and low-yielding samples respectively.

### Library preparation and RNA sequencing

Illumina libraries were prepared by Novogene Corporation Inc. (Chula Vista, CA) using the NEB Ultra II RNA Library Prep Kit for Illumina according to manufacturer protocols. The libraries were sequenced on an Illumina Novaseq 6000 S2 2 × 150 bp flow cell to an average depth of 29.24 × 10^6^ raw reads per sample for a total of 4.82^9^ reads. The sequencing provider filtered the reads to remove ones containing Illumina adaptors, or more than 10% indeterminate bases (“N”s), or having more than half bases with a phred score below 5. The remaining “clean” reads, representing 95.52% of raw reads, were processed further as described below.

### Alignment, quantification, and normalization

We used Salmon [30] to map the reads to the GRCm38 reference genome with the selective alignment strategy and to determine transcript abundances. The median-of-ratios method [31, 32] of DESeq2, which computes the ratio of each transcript’s abundance in each sample to the geometric mean of the expression across samples and then determines a size factor for each sample as the median of these ratios, was used to normalize gene expression to library size.

### Principal component analysis

Principal component analysis (PCA) was carried out with the prcomp R function with the scale parameter set to TRUE. Transcripts were filtered to remove those having a normalized count less than 5 in every sample—hereafter referred to as low-expressed transcripts.

### Hierarchical clustering and gene expression heatmaps

Samples or timepoints were clustered hierarchically and plotted with pheatmap on gene expression scaled between 0 and 1, after excluding the low-expressed transcripts, using Pearson correlation as a similarity metric.

### Outlier detection

Principal components analysis (PCA) was performed on the filtered data and the *z*-score of the *k*th principal component of each replicate *i* at a given time point was calculated as

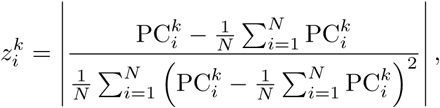

where *k* = 1, 2, 3, 4 are the top four PCs accounting for 96% of the total variation in the data and PC*^k^* is the principal component score of the *k*th component. A replicate was regarded as an outlier if *z^k^ >* 2 for any of the top four PCs. No outliers were detected in the data using this method.

In a complementary approach to identifying outliers we clustered all the samples hierarchically (Figs. S1 and S2). All three replicates for the 96h and 144h timepoints in IL3 conditions appeared to be visually significantly different from the time points immediately preceding or succeeding them. The time points immediately before and after the outliers were more correlated to each other than to the outliers (Fig. S3). These timepoints were unusual, having been preceded by supplementation with OHT (see above), so that the differences could be transient effects of OHT. We excluded these two timepoints from further analyses.

### Differential gene expression analysis

Differential expression analysis was conducted using DESeq2 [33], which uses the Wald test to determine the significance of the estimated fold change and the Benjamini-Hochberg procedure to correct for multiple testing. A transcript was considered differentially expressed if its adjusted *p*-value was less than 0.05 and the log_2_ fold change (FC) was more than 0.58 (±50% change).

### Non-negative matrix factorization

Non-negative matrix factorization (NMF) [24] was used to extract features, reduce dimensionality, and cluster the temporal expression patterns of all the transcripts in the dataset. Prior to NMF, the expression of each gene across all timepoints and conditions was scaled between 0 and 1.

#### NMF Implementation

The algorithm produces a lower-dimensional representation of the dataset by approximating the temporal pattern of each gene’s expression as a linear sum of a few characteristic temporal patterns, referred to here as *behaviors*. The algorithm does so by approximately factoring the gene expression matrix **X** into two lower-dimensional non-negative matrices, **W** and **H**, as

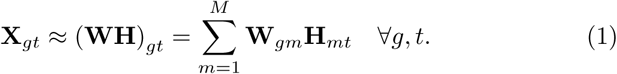

Here *g* = 1*, . . ., G*, *t* = 1*, . . ., T*, and *m* = 1*, . . ., M* index genes, timepoints, and behaviors respectively. *G*, *T*, and *M* are the total number of genes, timepoints, and behaviors respectively. **X***_gt_* is the expression of gene *g* at timepoint *t*. **H***_mt_* is the expression of behavior *m* and timepoint *t*. **W***_gm_* is the weight of behavior *m* in the linear combination of behaviors that approximates the expression pattern of gene *g*.

The algorithm iteratively updates the **W** and **H** matrices to converge to a local maximum of the objective function *O* given by

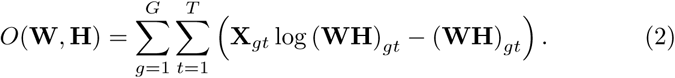

The alternating gradient method was used to produce a sequence {**W***^k^,* **H***^k^*}, where *k* = 1*, . . ., N*, by applying the multiplicative update rules

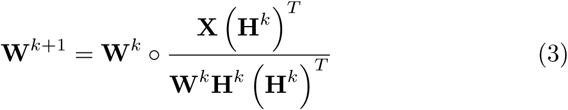

and

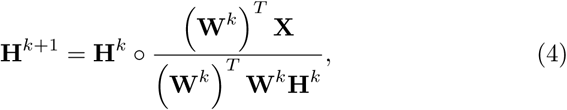

where ◦ signifies element-wise multiplication. The elements of the initial matrices *W* ^0^ and *H*^0^ were populated by sampling from the uniform distribution over [0, 1)

We calculated the Frobenius norm (*F^k^*) of the difference between original **X** matrix and its *k*th approximation, 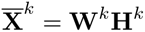,

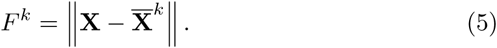

**W***^k^* and **H***^k^* were updated according to Equations 3 and 4 until the stopping criterion, that the absolute change in the Frobenius norm between successive iterations was less than the threshold *θ*,

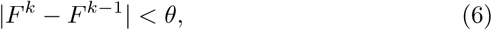

was met. Unless specified otherwise, *θ* was chosen to be 0.05.

#### Choice of the number of behaviors ***k***

The first local minimum of the root mean square error (RMSE),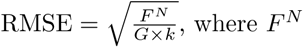, where *F^N^* is the Frobenius norm of the difference between the approximated and actual gene expression matrices, was found to occur at *k* = 7 (Fig. S4). We set the number of behaviors *k* = 10 to ensure that all potential behaviors had been recovered.

#### Robustness analysis

The robustness of the NMF algorithm to random initial conditions was checked by repeating the factorization 100 times and comparing the RMSE between each pair of approximated gene expression matrices to the RMSE between the approximated and actual gene expression matrices (Fig. S5). The RMSE between different approximations was found to be consistently lower than the RMSE between the approximated and actual gene expression matrices, demonstrating that the approximate solutions are robust to the random initialization of the NMF algorithm.

#### Visualization

The visualizations in Figure 5 were produced in two steps. In the first step, the behaviors were sorted according to the condition and time at which their expression peaked so that behaviors peaking earliest in GCSF conditions had the lowest rank, followed by those expressed later in GCSF, then by those expressed early in IL3, and finally by those expressed late in IL3. In the second step, for each gene, the behavior with the highest weight—the dominant behavior—was identified. The genes were grouped according to their dominant behavior and then sorted in descending order of the weight of the dominant behavior.

### Gene ontology analysis

`clusterProfiler` [34] was used to identify over-represented biological process Gene Ontology (GO) terms associated with differentially expressed genes (DEGs) [35] or the top 500 genes associated with a behavior, and having a maximum normalized expression of at least 3, using all annotated genes as the background gene list. A functional annotation category was considered significant if the Benjamini-Hochberg adjusted *p*-value was less than 0.05.

## Supporting information

**S1 Fig.**
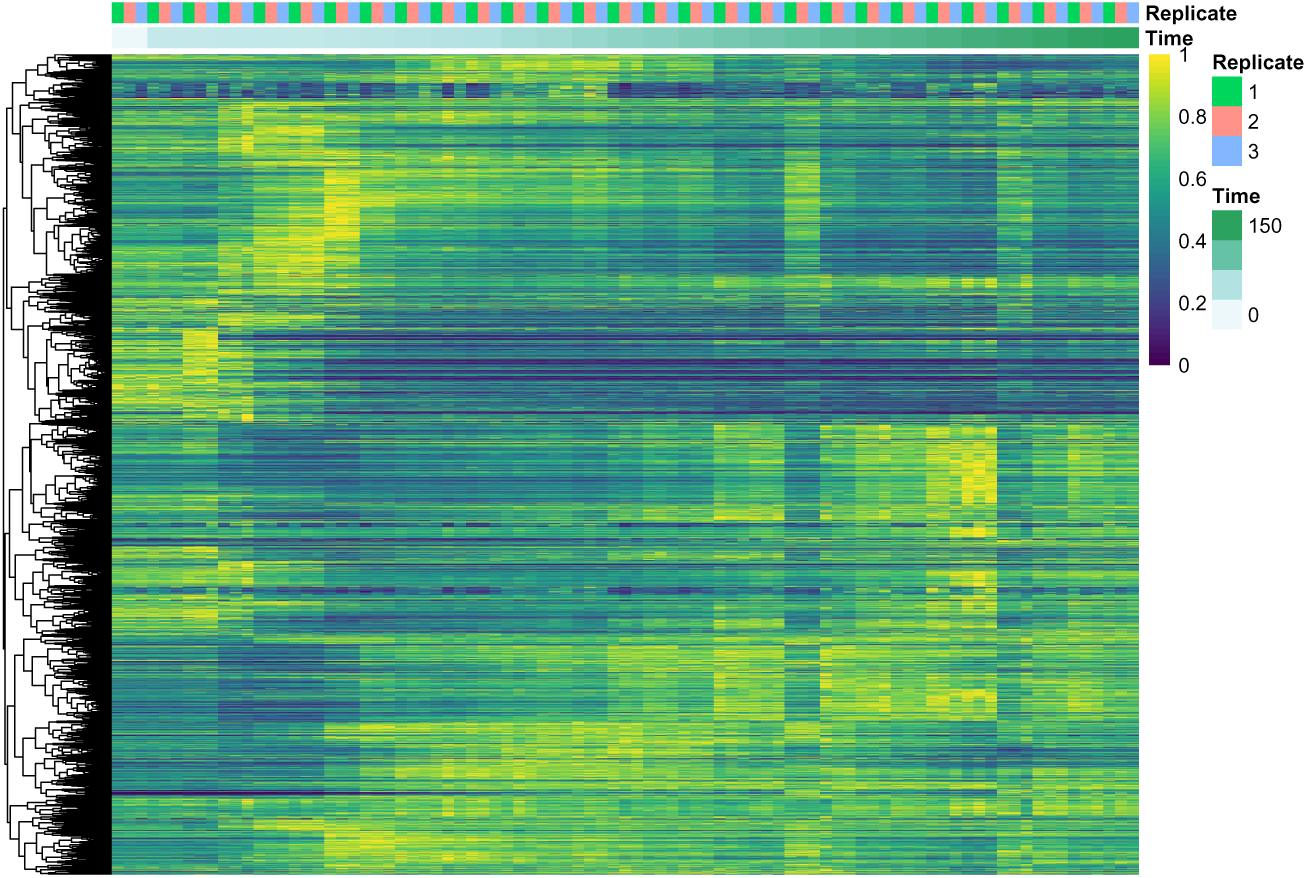
Genome-wide gene expression in all IL3 samples. Color indicates the expression of each transcript (*y*-axis) in each IL3 sample (*x*-axis) scaled from 0 to 1. Transcripts were clustered hierarchically using Pearson correlation as a similarity measure.

**S2 Fig.**
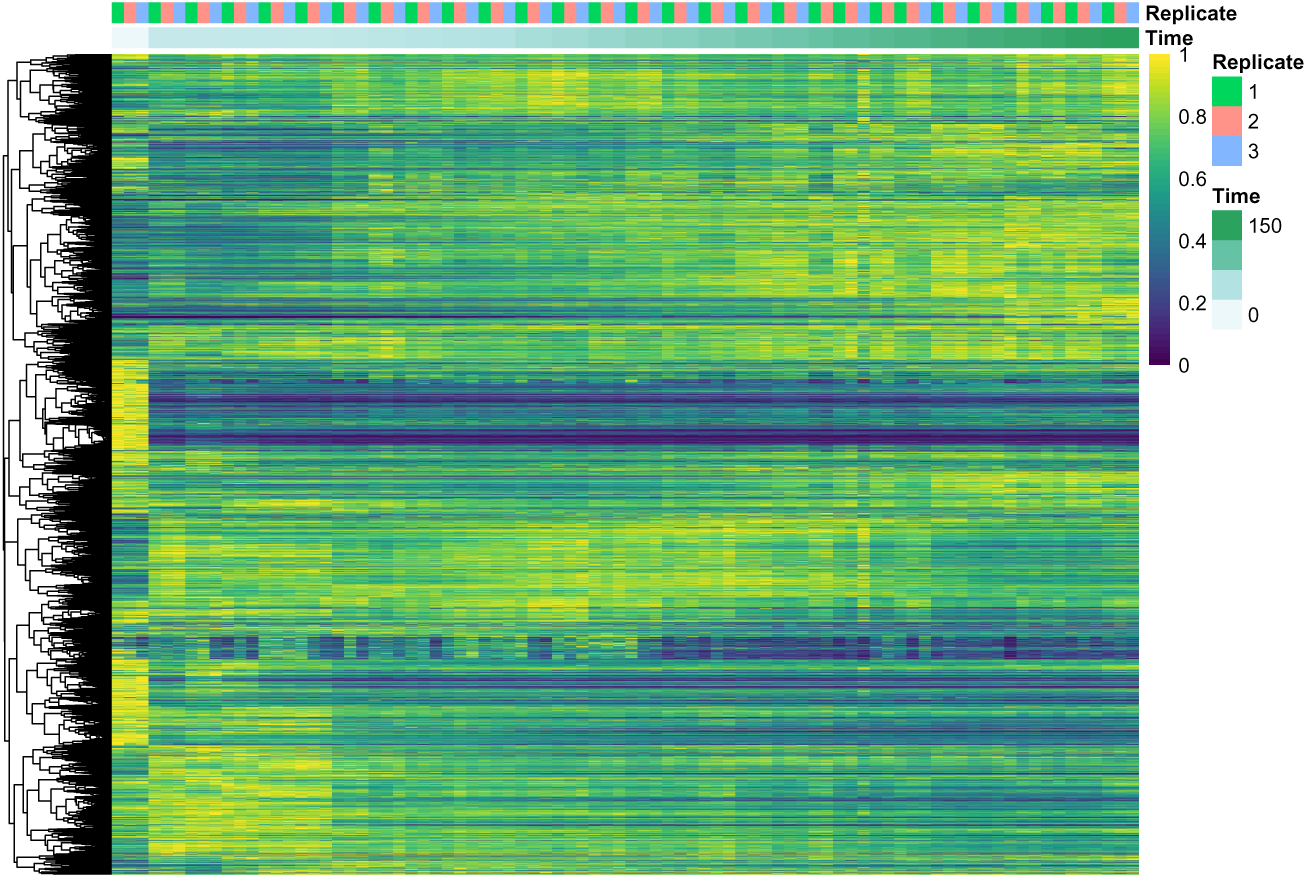
Genome-wide gene expression in all GCSF samples. Color indicates the expression of each transcript (*y*-axis) in each GCSF sample (*x*-axis) scaled from 0 to 1. Transcripts were clustered hierarchically using Pearson correlation as a similarity measure.

**S3 Fig.**
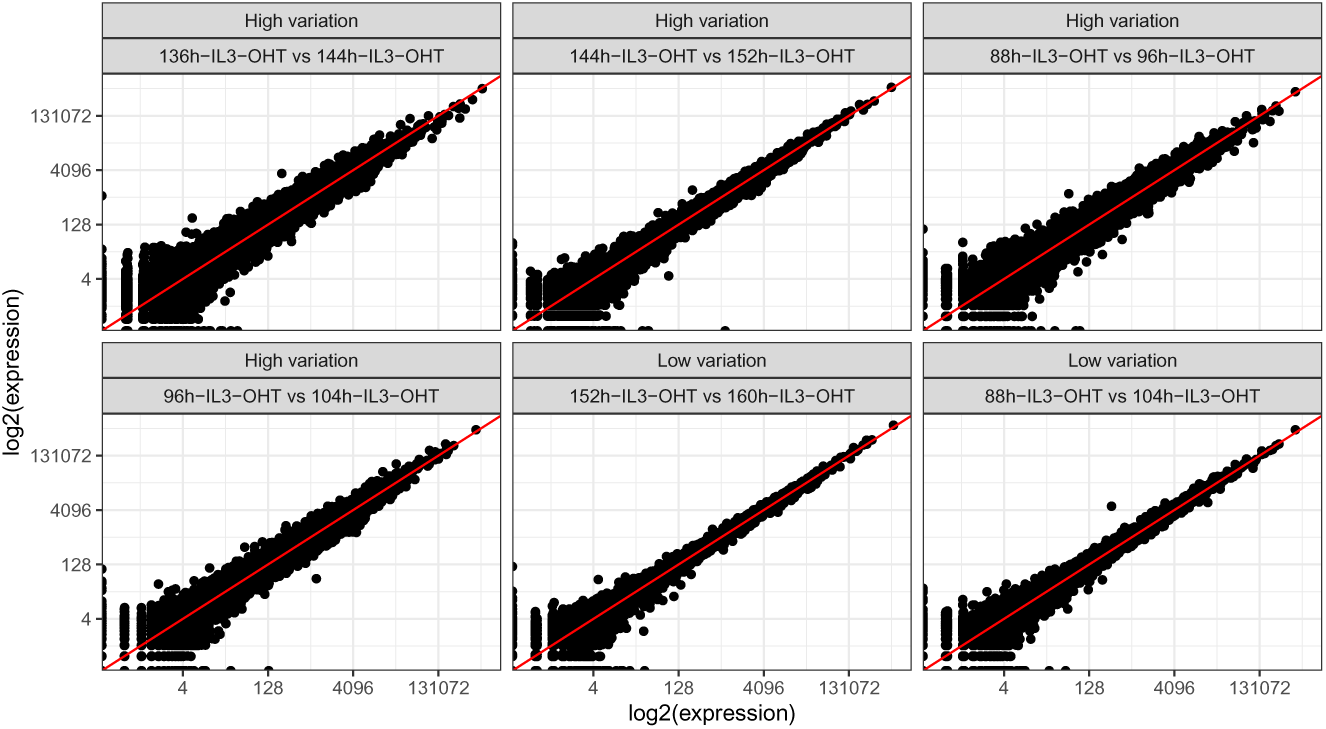
Correlation in gene expression between selected samples. Correlation in gene expression between suspected outliers and immediate neighbors (88h vs 96h, 96h vs 104h, 136h vs 144h, and 144h vs 152h). Correlation between the timepoint immediately preceding and the timepoint immediately following the suspected outlier (88h vs 104h and 136h vs 152h).

**S4 Fig.**
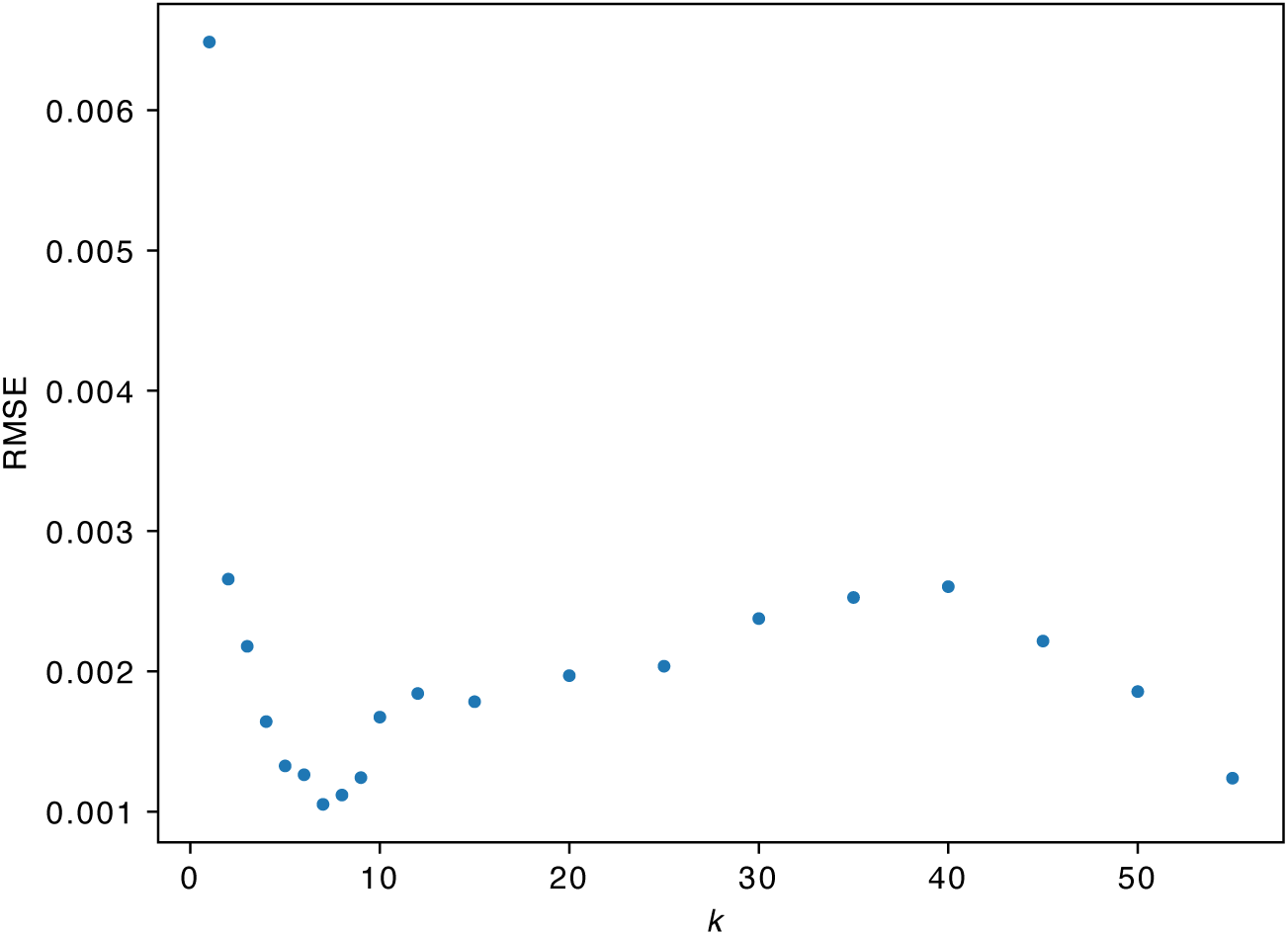
The error of the NMF approximation vs. the number of metagenes. The root mean square error (RMSE) between the gene expression matrix and its NMF approximation as the number of metagenes (*k*) is varied.

**S5 Fig.**
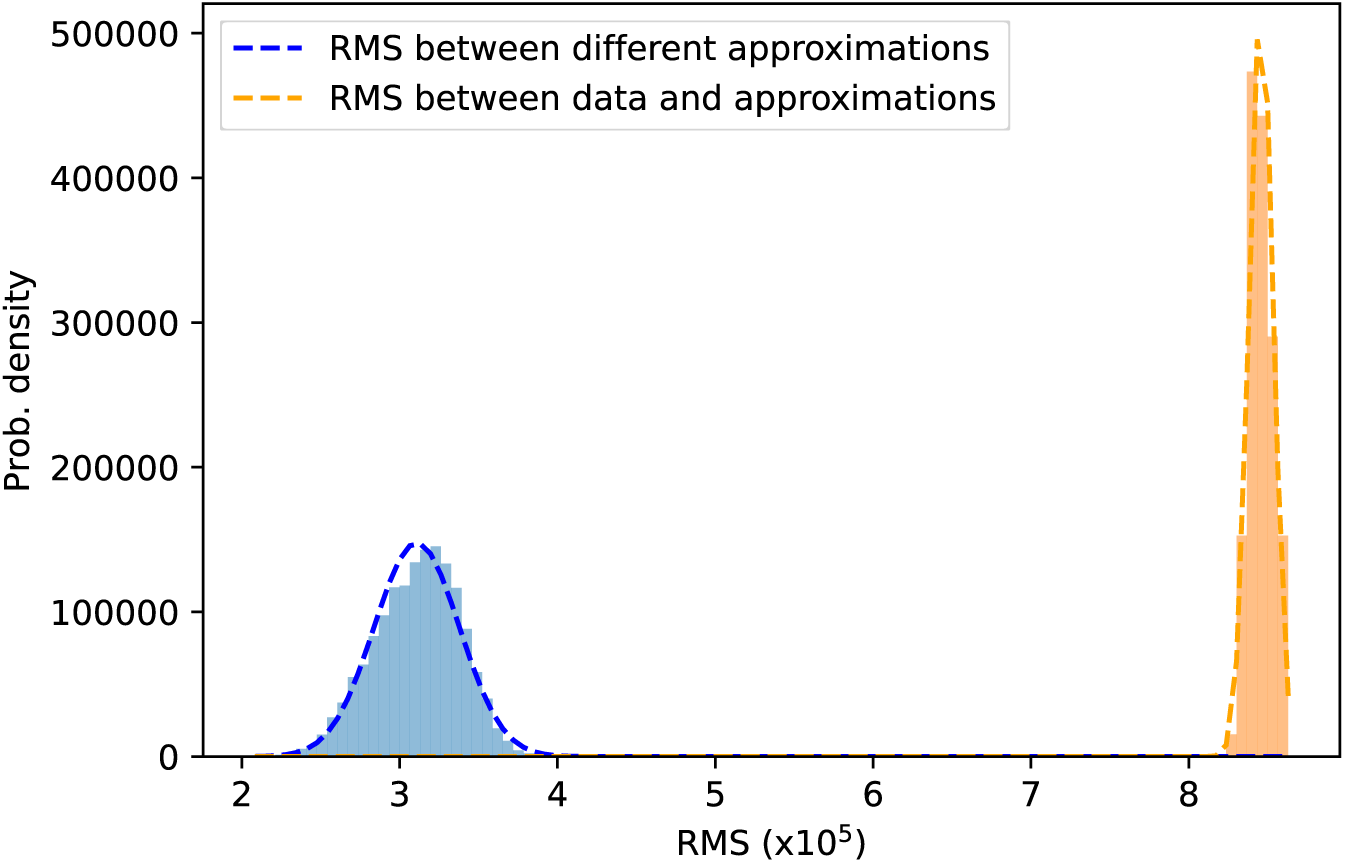
Robustness of NMF. Histograms of the RMSE between the gene expression matrix and 100 NMF approximations (orange) or the RMSE between the approximations (blue). Dashed lines are the Gaussian densities with the mean and standard deviation of each histogram.

**S6 Fig.**
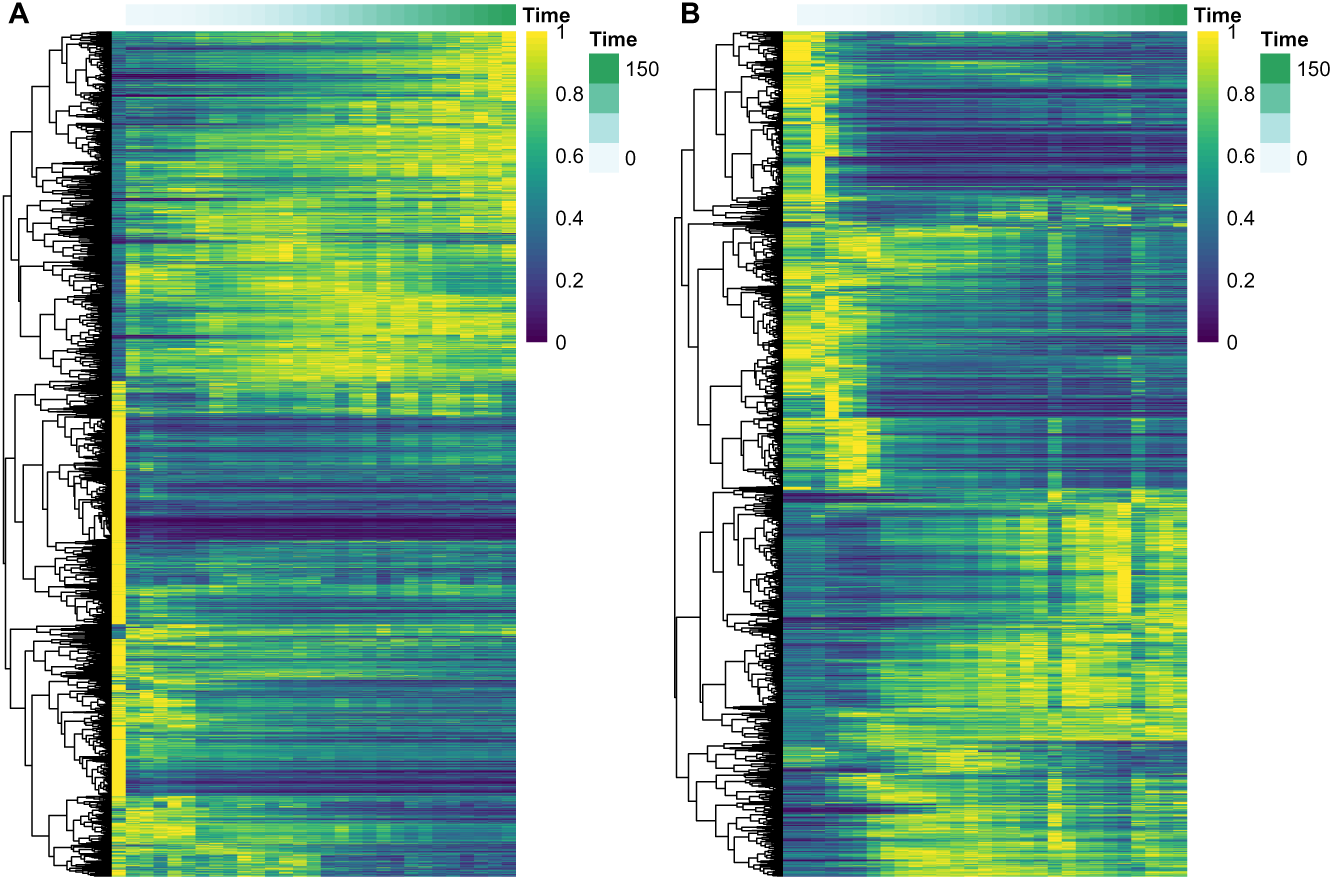
Transient expression of differentially expressed genes. The expression of all differentially expressed genes, scaled between 0 and 1, is shown as a color map. Genes are clustered hierarchically according to the Pearson correlation of scaled temporal expression patterns. (A) GCSF. Plotted genes are the union of DEGs identified in the comparisons between the −48*h* and 168*h*, −48*h* and 0*h*, and 0*h* and 168*h* GCSF samples (Fig. 3A–C). (B) IL3. DEGs identified in the comparison between the 0*h* and 168*h* IL3 samples (Fig. 3D).

**S7 Fig.**
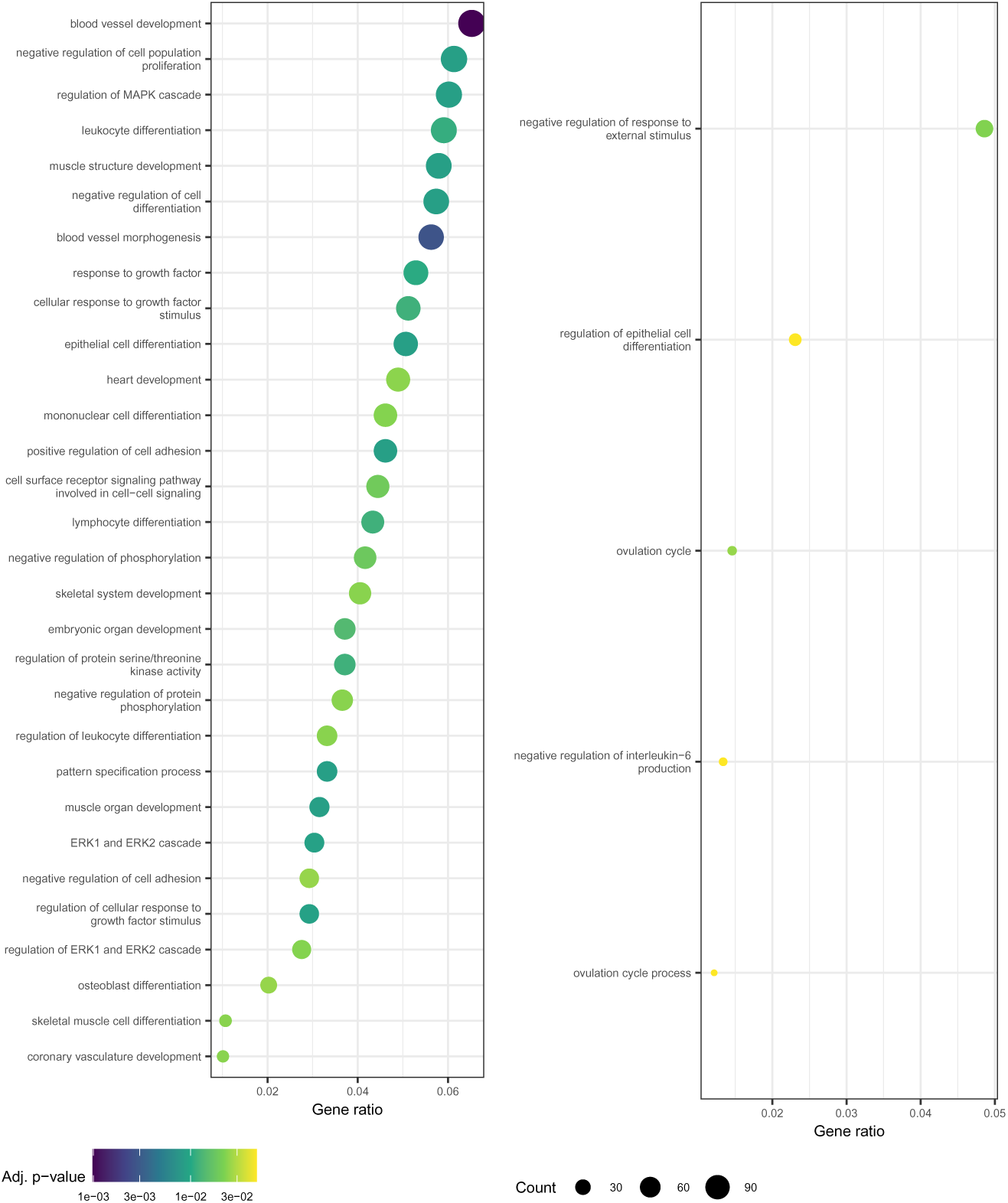
GO terms enriched for genes differentially expressed between. 1**h and** 2**h and** 2**h and** 3**h IL3 samples.** GO terms enriched for DEGs between 1h and 2h and between 2h and 3h are shown on the left and right respectively. See the legend of Fig. 7 for plot description.

**S8 Fig.**
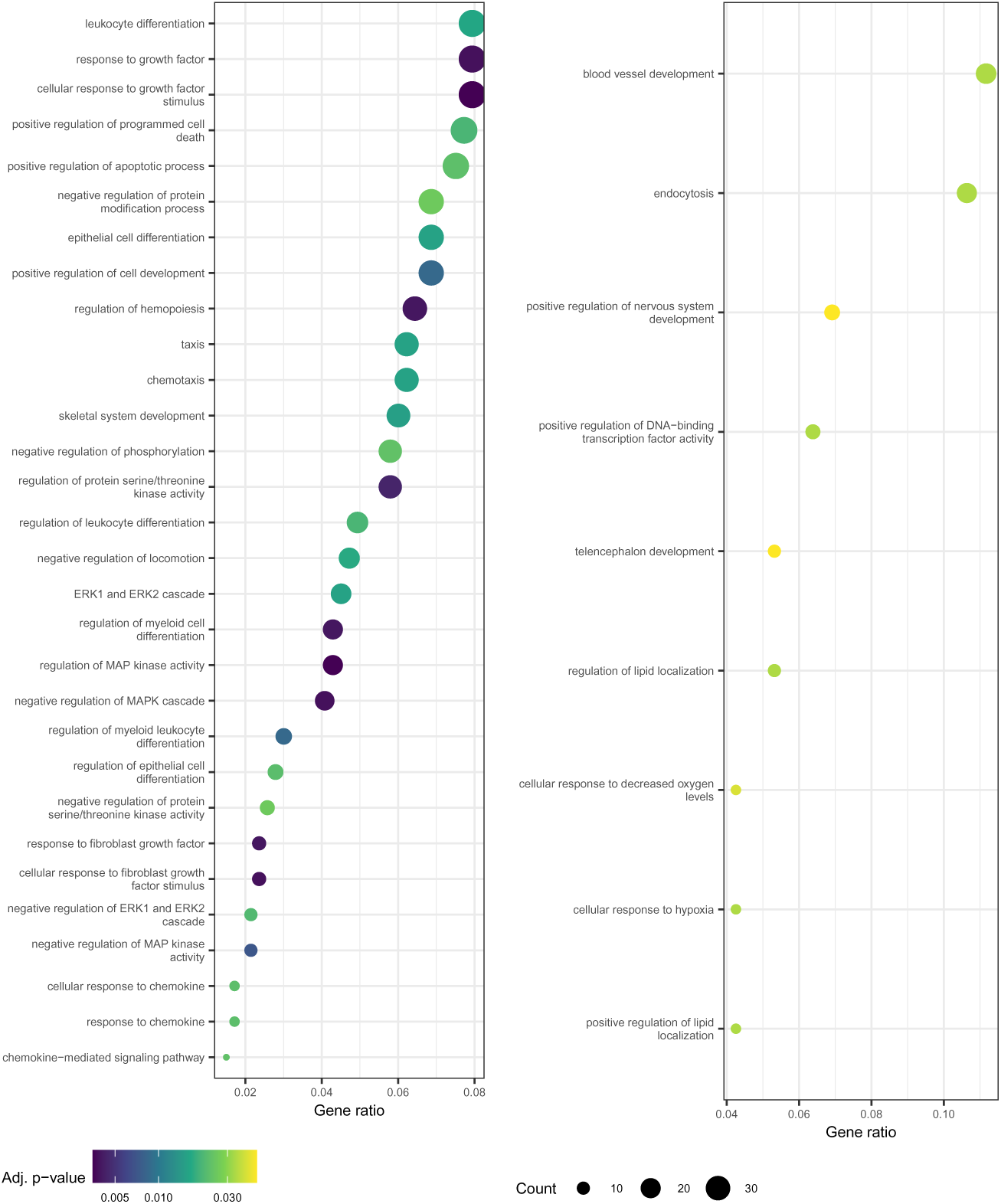
GO terms enriched for genes differentially expressed between. 1**h and** 2**h and** 2**h and** 3**h GCSF samples.** GO terms enriched for DEGs between 1h and 2h and between 2h and 3h are shown on the left and right respectively. See the legend of Fig. 7 for plot description.

**S9 Fig.**
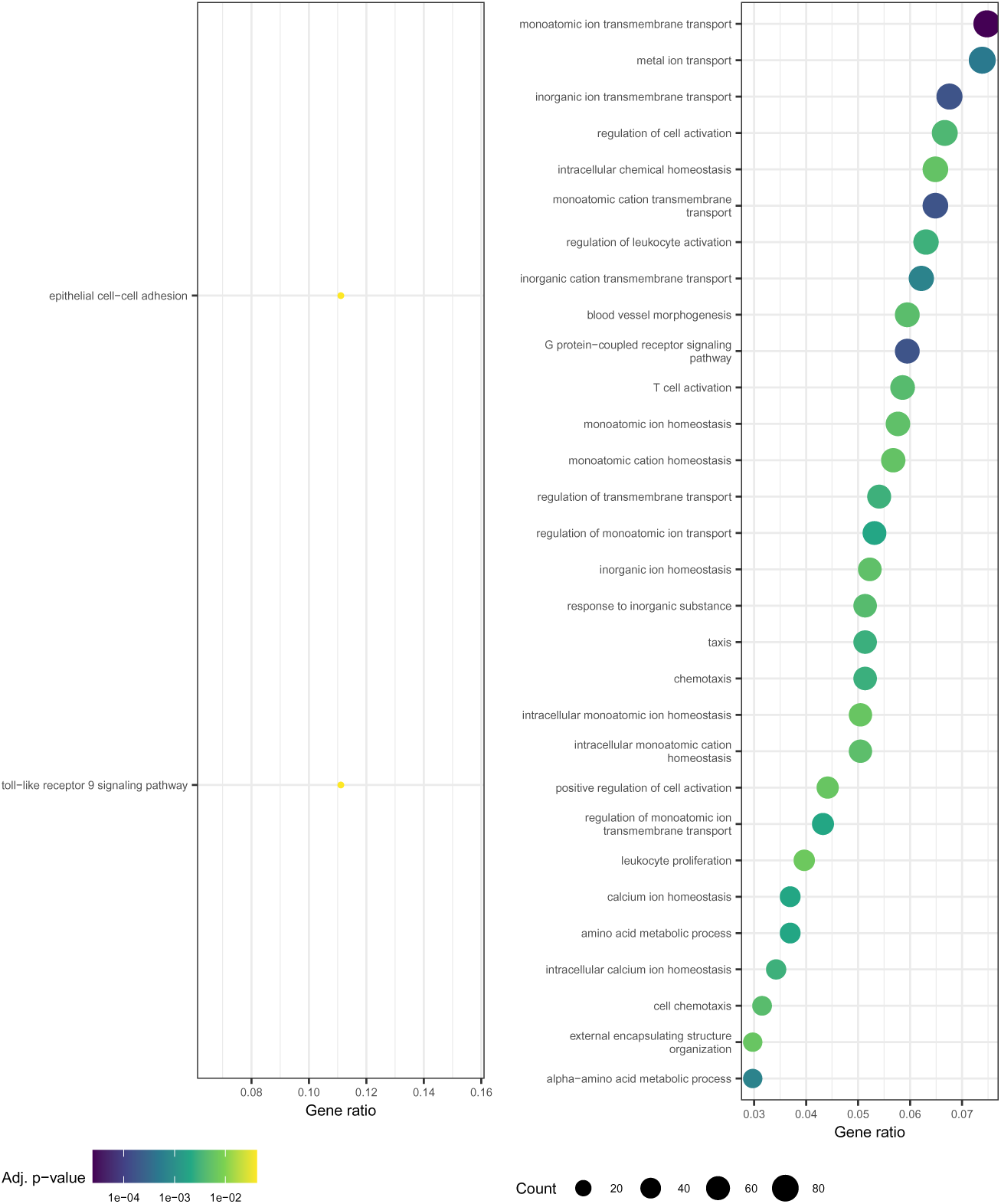
GO terms enriched for genes differentially expressed between. 3**h and** 4**h and** 4**h and** 8**h IL3 samples.** GO terms enriched for DEGs between 3h and 4h and between 4h and 8h are shown on the left and right respectively. See the legend of Fig. 7 for plot description.

**S10 Fig.**
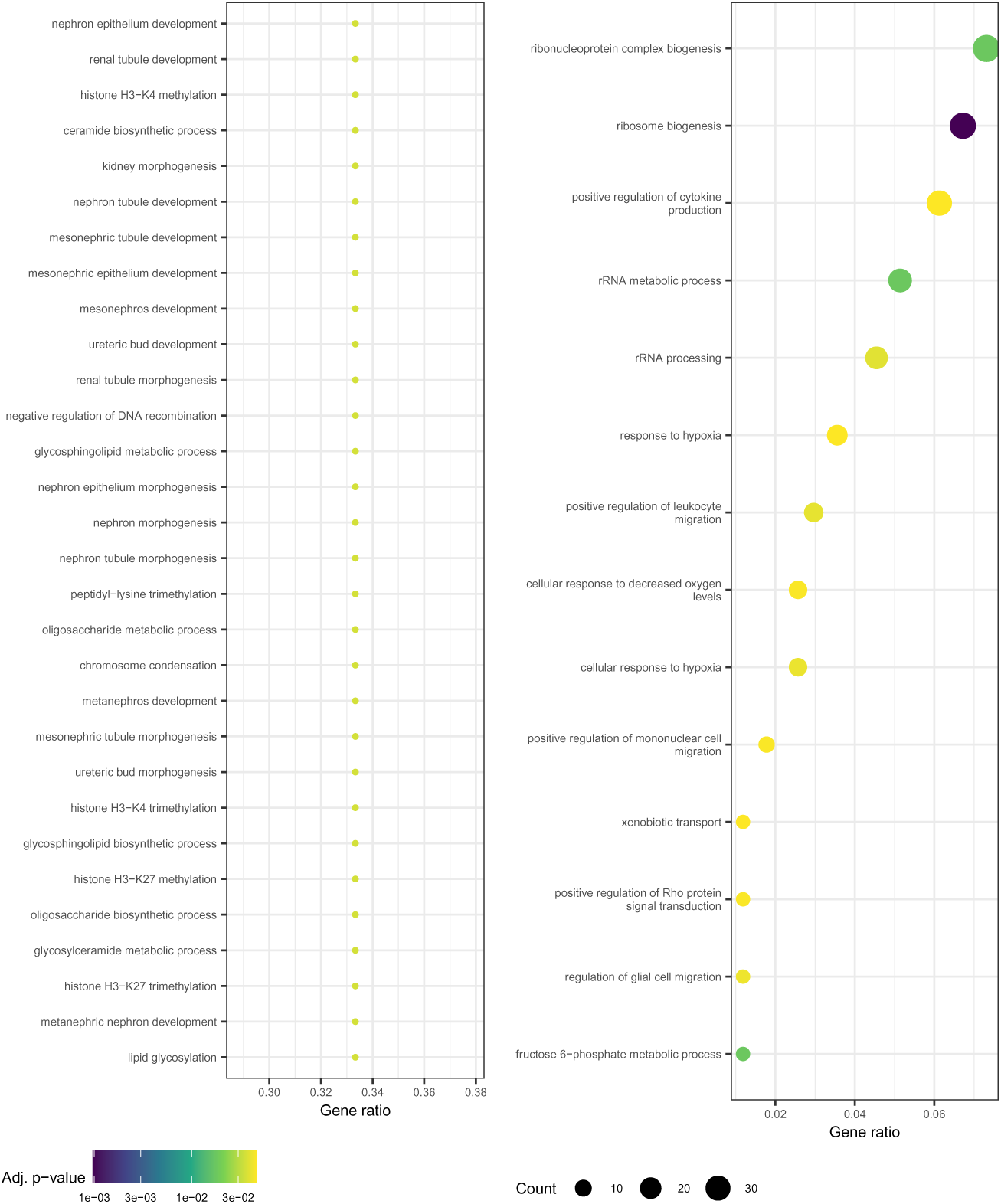
GO terms enriched for genes differentially expressed between. 3**h and** 4**h and** 4**h and** 8**h GCSF samples.** GO terms enriched for DEGs between 3h and 4h and between 4h and 8h are shown on the left and right respectively. See the legend of Fig. 7 for plot description.

**S11 Fig.**
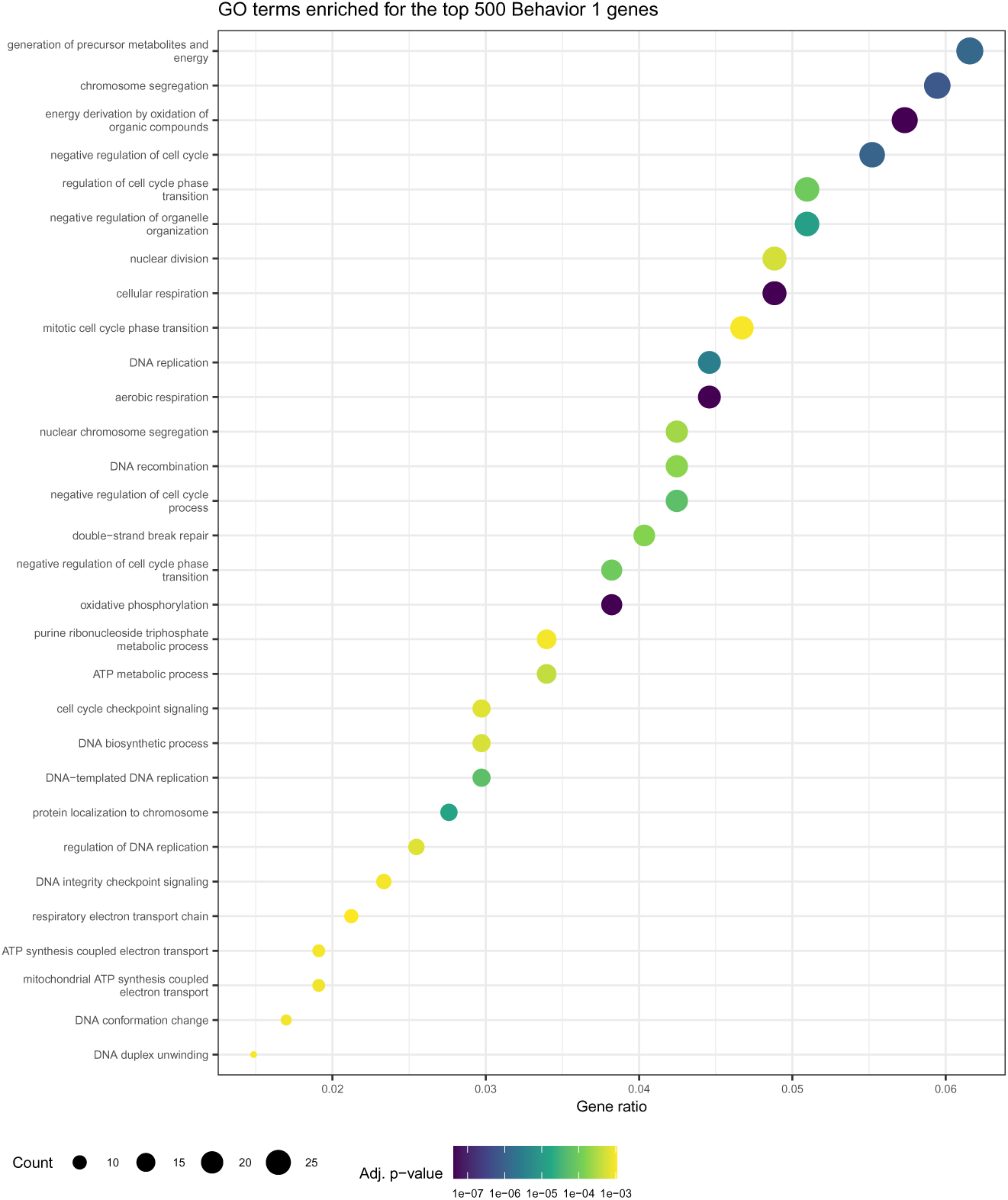
GO terms enriched for the top 500 behavior 1 genes. See the legend of Fig. 7 for plot description.

**S12 Fig.**
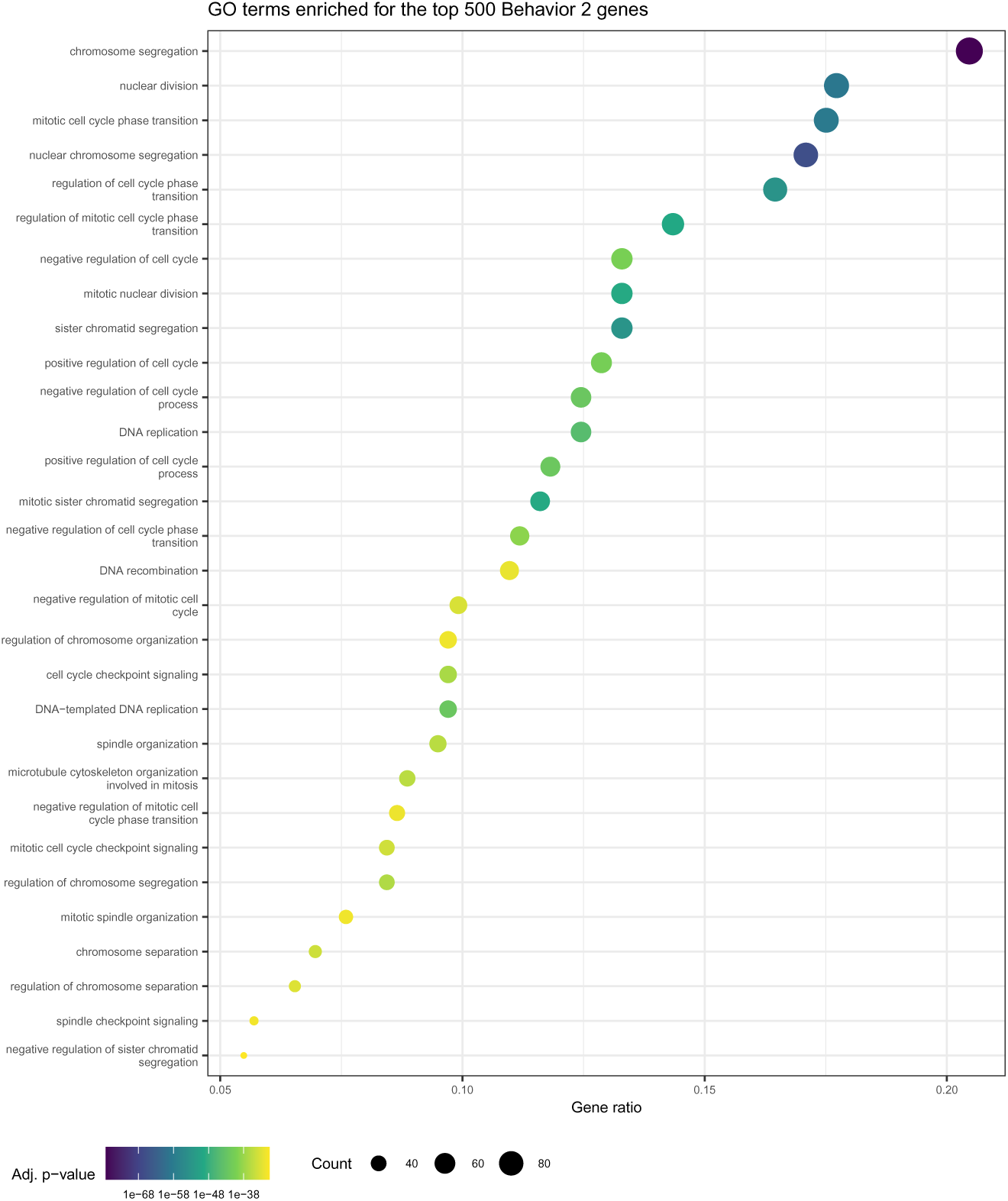
GO terms enriched for the top 500 behavior 2 genes. See the legend of Fig. 7 for plot description.

**S13 Fig.**
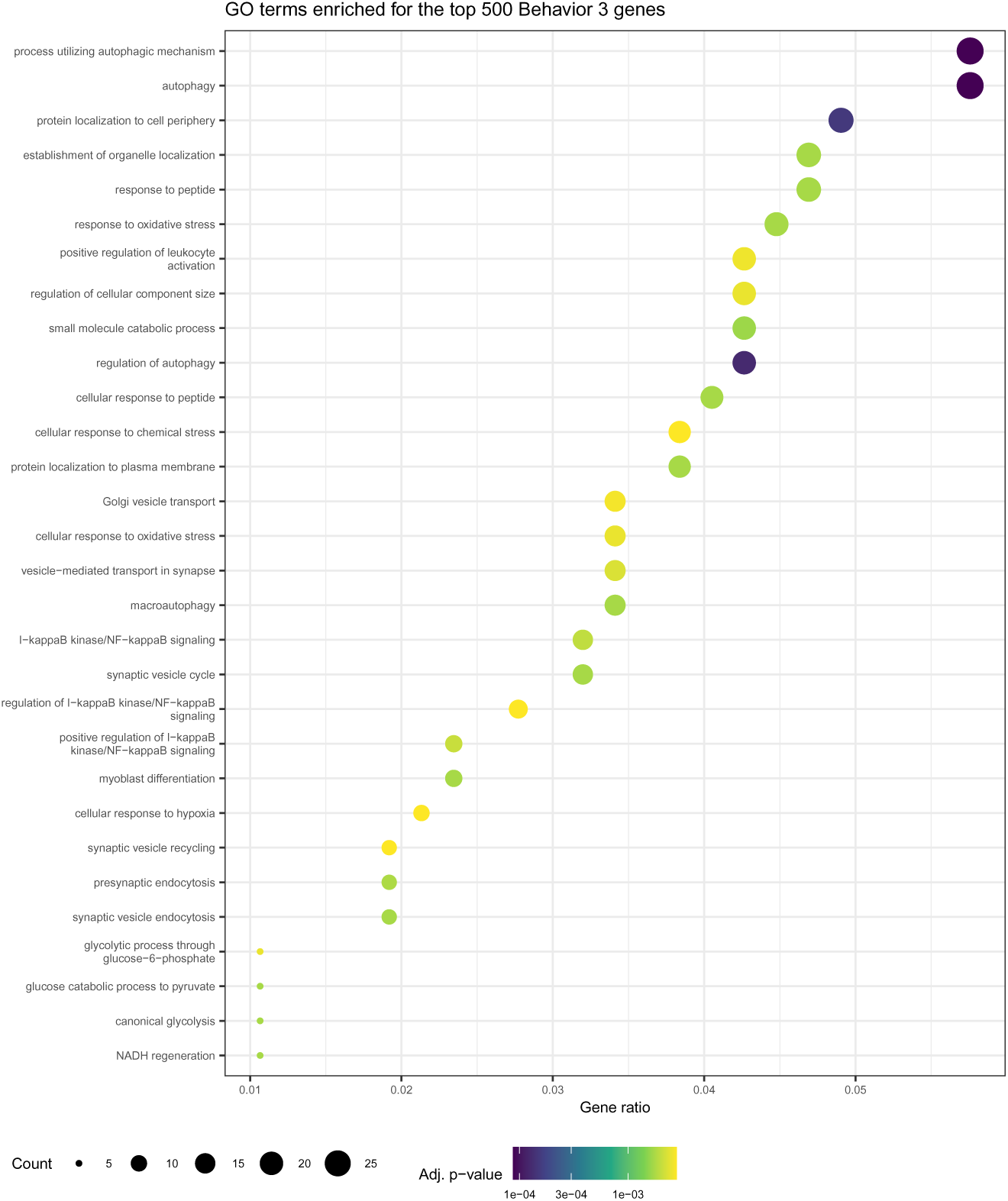
GO terms enriched for the top 500 behavior 3 genes. See the legend of Fig. 7 for plot description.

**S14 Fig.**
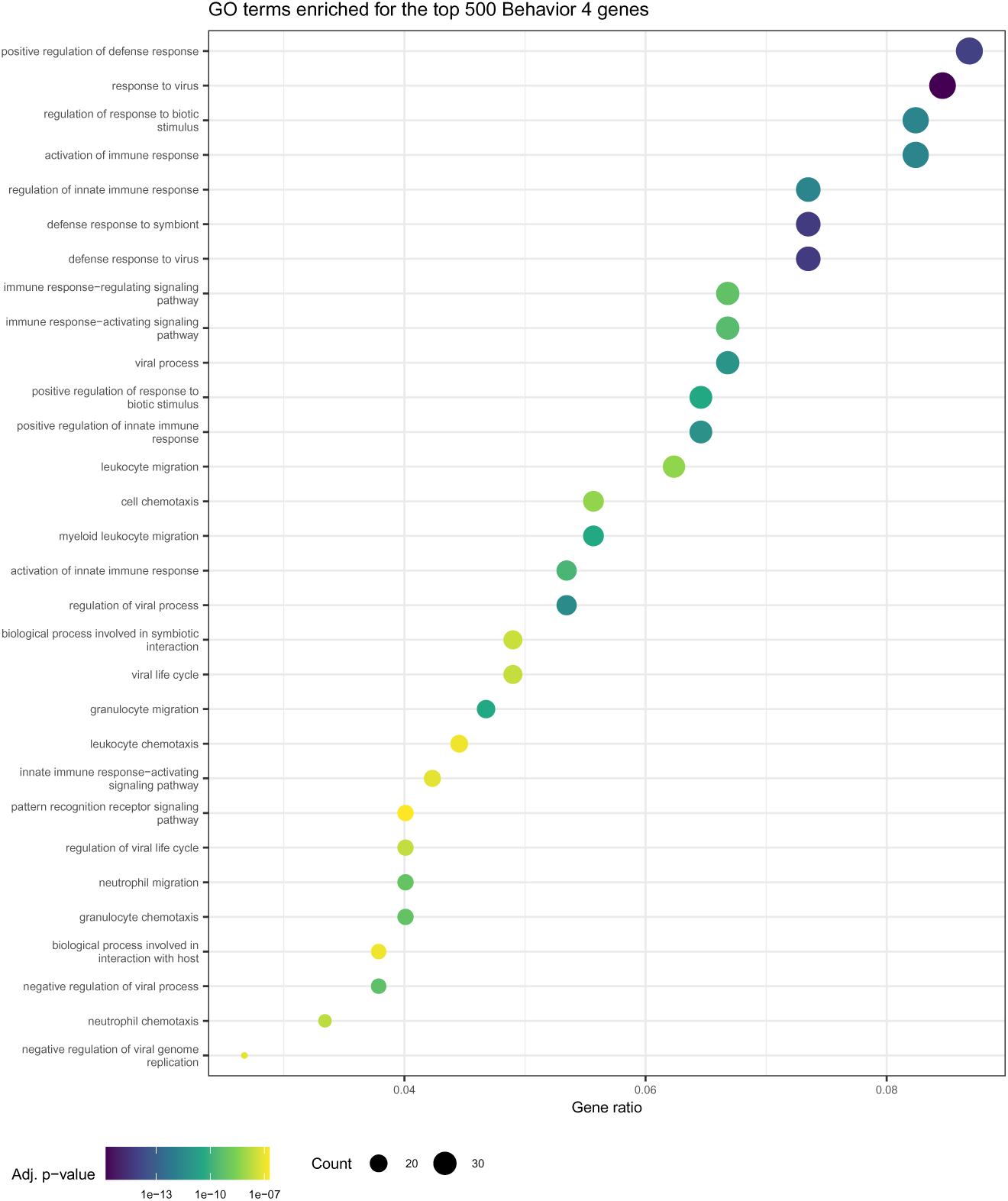
GO terms enriched for the top 500 behavior 4 genes. See the legend of Fig. 7 for plot description.

**S15 Fig.**
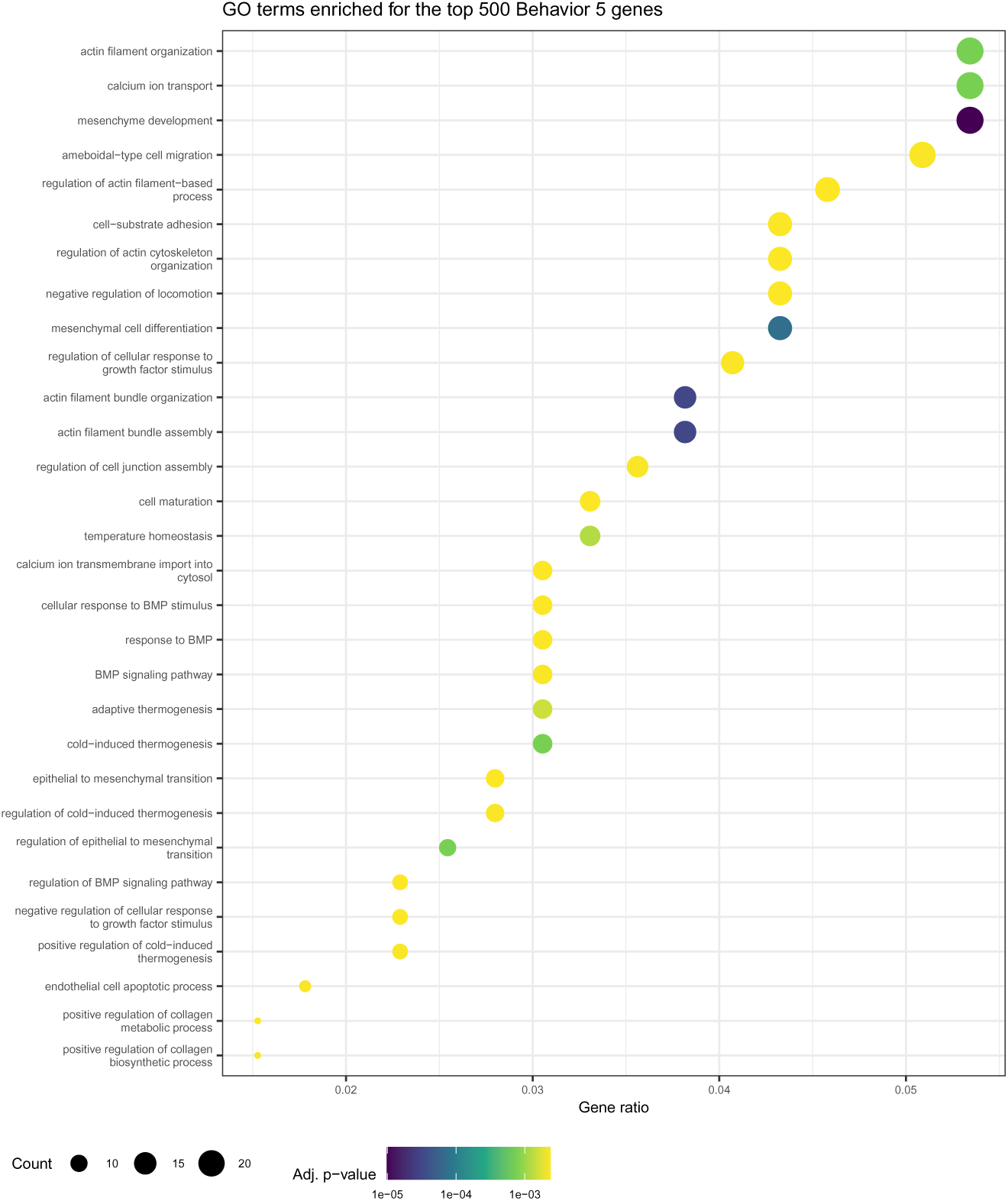
GO terms enriched for the top 500 behavior 5 genes. See the legend of Fig. 7 for plot description.

**S16 Fig.**
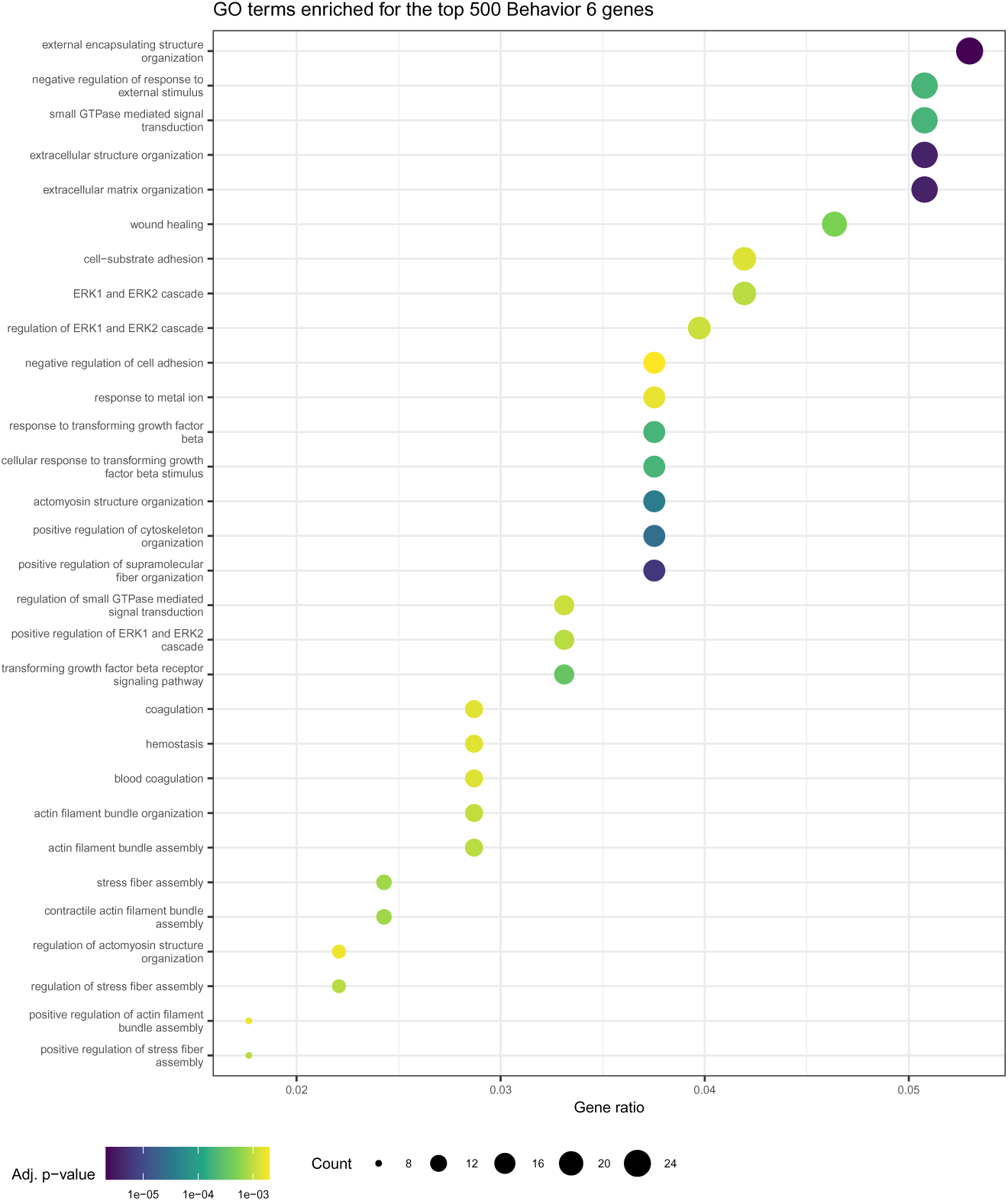
GO terms enriched for the top 500 behavior 6 genes. See the legend of Fig. 7 for plot description.

**S17 Fig.**
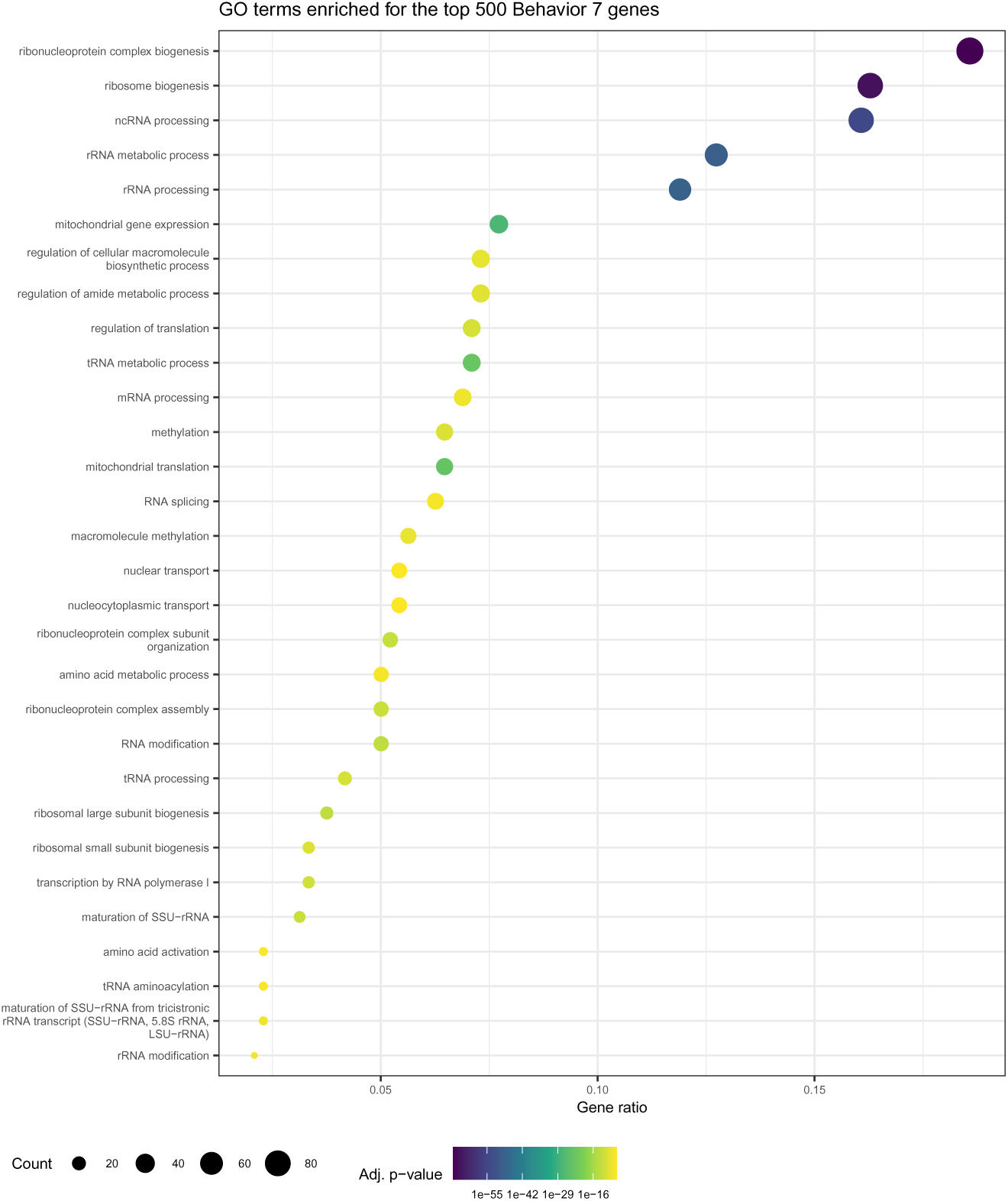
GO terms enriched for the top 500 behavior 7 genes. See the legend of Fig. 7 for plot description.

**S18 Fig.**
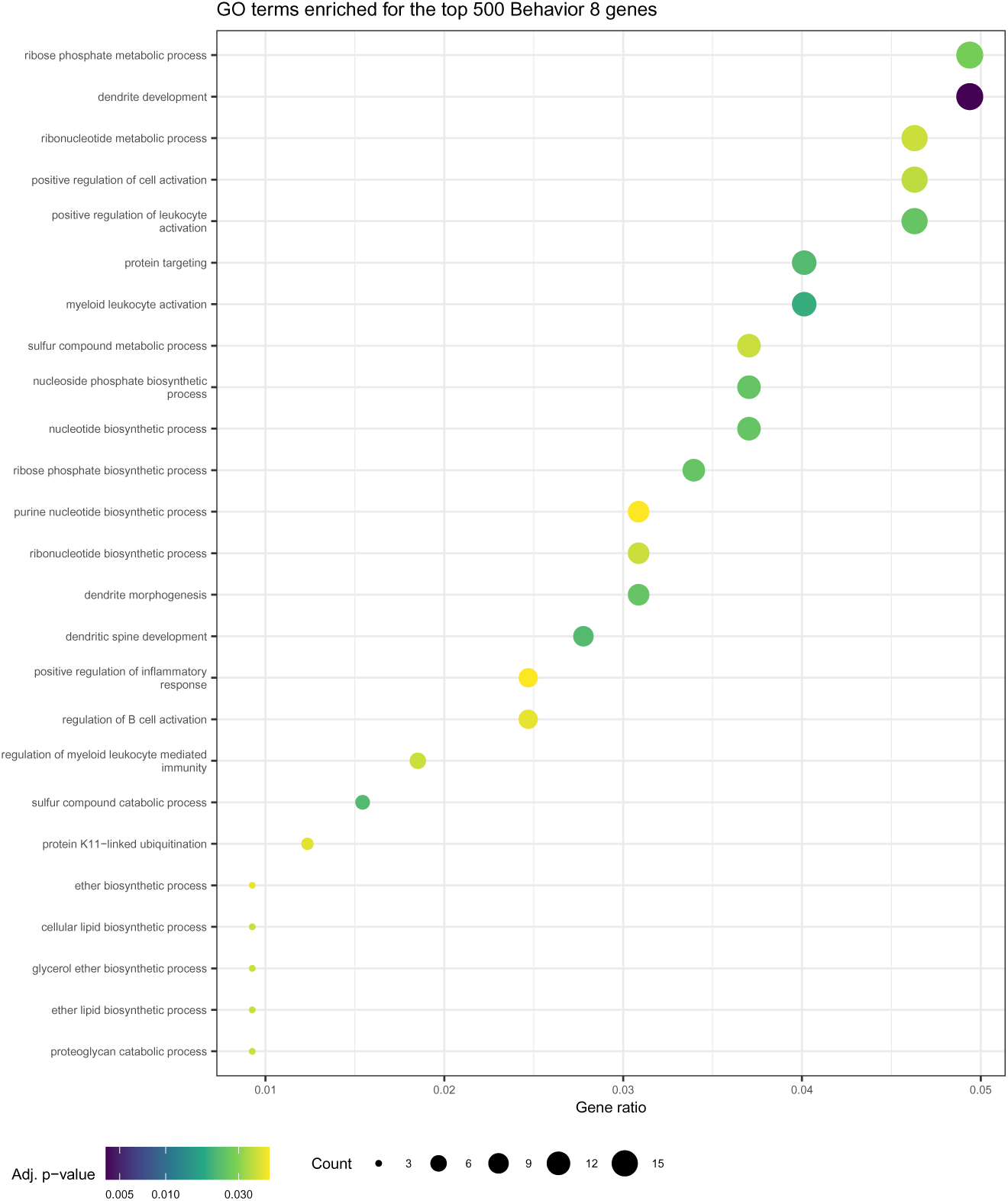
GO terms enriched for the top 500 behavior 8 genes. See the legend of Fig. 7 for plot description.

**S19 Fig.**
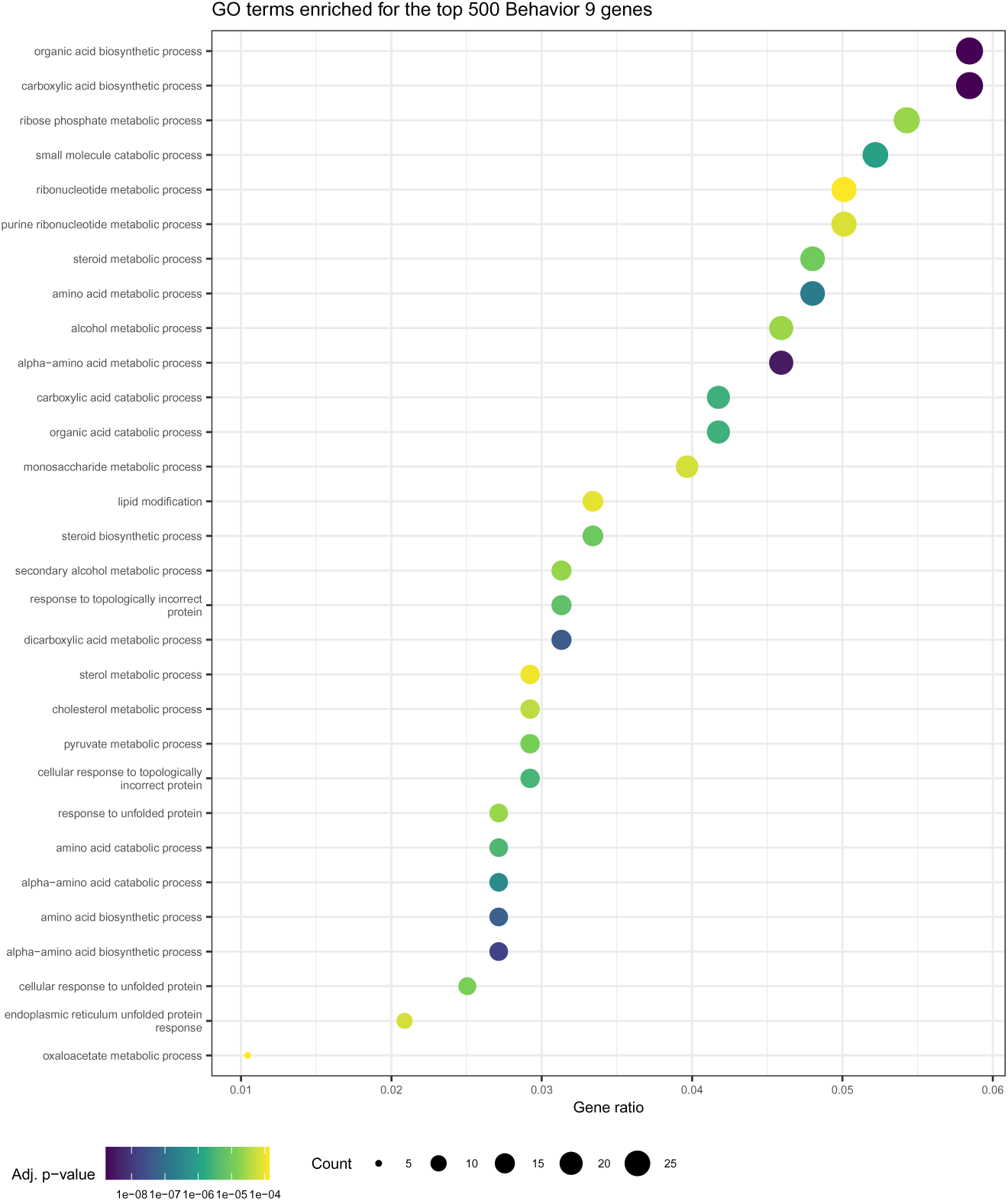
GO terms enriched for the top 500 behavior 9 genes. See the legend of Fig. 7 for plot description.

**S20 Fig.**
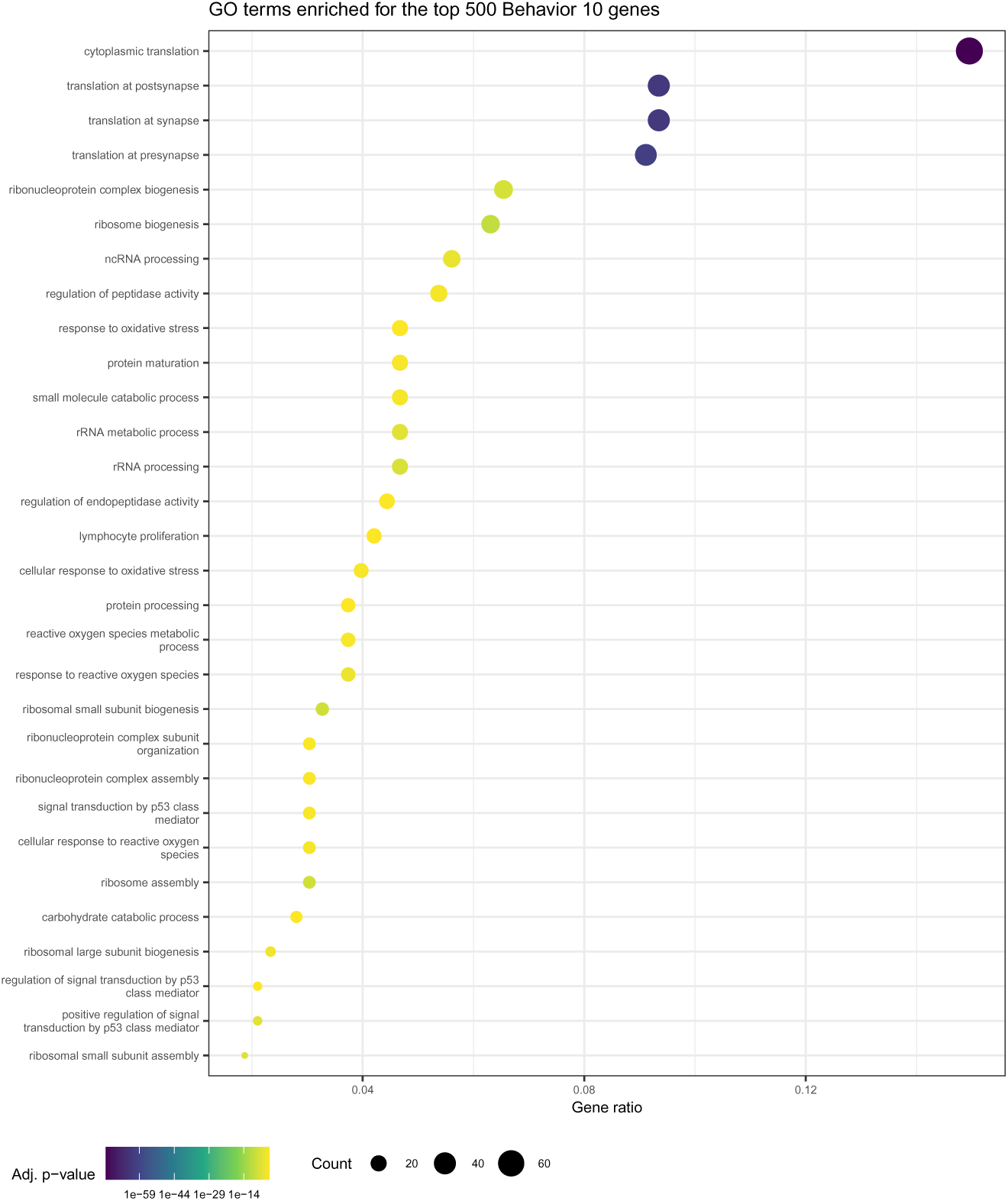
GO terms enriched for the top 500 behavior 10 genes. See the legend of Fig. 7 for plot description.

## Acknowledgments

We thank the UND Genomics core for services. This work was supported by the National Science Foundation [1942471 to M.].

## References

1. Görgens A, Radtke S, Horn PA, Giebel B. New relationships of human hematopoietic lineages facilitate detection of multipotent hematopoietic stem and progenitor cells. Cell cycle (Georgetown, Tex). 2013;12(22):3478–3482. doi:10.4161/cc.26900.

2. Keohane E. Rodak’s hematology : clinical principles and applications. St. Louis, Missouri: Elsevier; 2020.

3. Laslo P, Spooner CJ, Warmflash A, Lancki DW, Lee HJ, Sciammas R, et al. Multilineage transcriptional priming and determination of alternate hematopoietic cell fates. Cell. 2006;126(4):755–66. doi:10.1016/j.cell.2006.06.052.

4. Olsson A, Venkatasubramanian M, Chaudhri VK, Aronow BJ, Salomonis N, Singh H, et al. Single-cell analysis of mixed-lineage states leading to a binary cell fate choice. Nature. 2016;537(7622):698–702. doi:10.1038/nature19348.

5. Scott EW, Simon MC, Anastasi J, Singh H. Requirement of transcription factor PU.1 in the development of multiple hematopoietic lineages. Science. 1994;265(5178):1573–7.

6. Zhang DE, Zhang P, Wang ND, Hetherington CJ, Darlington GJ, Tenen DG. Absence of granulocyte colony-stimulating factor signaling and neutrophil development in CCAAT enhancer binding protein alpha-deficient mice. Proc Natl Acad Sci U S A. 1997;94(2):569–74.

7. Dahl R, Walsh JC, Lancki D, Laslo P, Iyer SR, Singh H, et al. Regulation of macrophage and neutrophil cell fates by the PU.1:C/EBPalpha ratio and granulocyte colony-stimulating factor. Nat Immunol. 2003;4(10):1029–36. doi:10.1038/ni973.

8. Olsson A, Venkatasubramanian M, Chaudhri VK, Aronow BJ, Salomonis N, Singh H, et al. Single-cell analysis of mixed-lineage states leading to a binary cell fate choice. Nature. 2016;537(7622):698–702. doi:10.1038/nature19348.

9. Novershtern N, Subramanian A, Lawton LN, Mak RH, Haining WN, McConkey ME, et al. Densely interconnected transcriptional circuits control cell states in human hematopoiesis. Cell. 2011;144(2):296–309. doi:10.1016/j.cell.2011.01.004.

10. May G, Soneji S, Tipping AJ, Teles J, McGowan SJ, Wu M, et al. Dynamic analysis of gene expression and genome-wide transcription factor binding during lineage specification of multipotent progenitors. Cell Stem Cell. 2013;13(6):754–68. doi:10.1016/j.stem.2013.09.003.

11. Xie H, Ye M, Feng R, Graf T. Stepwise reprogramming of B cells into macrophages. Cell. 2004;117(5):663–76.

12. Keightley MC, Carradice DP, Layton JE, Pase L, Bertrand JY, Wittig JG, et al. The Pu.1 target gene Zbtb11 regulates neutrophil development through its integrase-like HHCC zinc finger. Nature Communications. 2017;8(1):14911. doi:10.1038/ncomms14911.

13. Guanglan L, Wenke H, Wenxue H. Transcription factor PU.1 and immune cell differentiation (Review). Int J Mol Med. 2020;46(6):1943–1950.

14. Bertolino E, Reinitz J, Manu. The analysis of novel distal Cebpa enhancers and silencers using a transcriptional model reveals the complex regulatory logic of hematopoietic lineage specification. Dev Biol. 2016;413(1):128–44. doi:10.1016/j.ydbio.2016.02.030.

15. Repele A, Krueger S, Bhattacharyya T, Tuineau MY, Manu. The regulatory control of Cebpa enhancers and silencers in the myeloid and red-blood cell lineages. PLoS One. 2019;14(6):e0217580. doi:10.1371/journal.pone.0217580.

16. Walsh JC, DeKoter RP, Lee HJ, Smith ED, Lancki DW, Gurish MF, et al. Cooperative and antagonistic interplay between PU.1 and GATA-2 in the specification of myeloid cell fates. Immunity. 2002;17(5):665–76.

17. Tusi BK, Wolock SL, Weinreb C, Hwang Y, Hidalgo D, Zilionis R, et al. Population snapshots predict early haematopoietic and erythroid hierarchies. Nature. 2018;555(7694):54–60. doi:10.1038/nature25741.

18. Haghverdi L, Lun ATL, Morgan MD, Marioni JC. Batch effects in single-cell RNA-sequencing data are corrected by matching mutual nearest neighbors. Nat Biotechnol. 2018;36(5):421–427. doi:10.1038/nbt.4091.

19. Weinreb C, Wolock S, Tusi BK, Socolovsky M, Klein AM. Fundamental limits on dynamic inference from single-cell snapshots. Proceedings of the National Academy of Sciences. 2018;115(10):E2467–E2476. doi:10.1073/pnas.1714723115.

20. Weinreb C, Wolock S, Klein AM. SPRING: a kinetic interface for visualizing high dimensional single-cell expression data. Bioinformatics. 2017;34(7):1246–1248. doi:10.1093/bioinformatics/btx792.

21. Weinreb C, Wolock S, Klein AM. SPRING: a kinetic interface for visualizing high dimensional single-cell expression data. Bioinformatics. 2018;34(7):1246–1248. doi:10.1093/bioinformatics/btx792.

22. Franzke A. The role of G-CSF in adaptive immunity. Cytokine Growth Factor Rev. 2006;17(4):235–244.

23. Lee DD, Seung HS. Learning the parts of objects by non-negative matrix factorization. Nature. 1999;401(6755):788–91. doi:10.1038/44565.

24. Brunet JP, Tamayo P, Golub TR, Mesirov JP. Metagenes and molecular pattern discovery using matrix factorization. Proc Natl Acad Sci U S A. 2004;101(12):4164–9. doi:10.1073/pnas.0308531101.

25. Handzlik JE, Manu. Data-driven modeling predicts gene regulatory network dynamics during the differentiation of multipotential hematopoietic progenitors. PLOS Computational Biology. 2022;18(1):1–31. doi:10.1371/journal.pcbi.1009779.

26. Choi J, Lysakovskaia K, Stik G, Demel C, Soding J, Tian TV, et al. Evidence for additive and synergistic action of mammalian enhancers during cell fate determination. eLife. 2021;10:e65381. doi:10.7554/eLife.65381.

27. Buenrostro JD, Giresi PG, Zaba LC, Chang HY, Greenleaf WJ. Transposition of native chromatin for fast and sensitive epigenomic profiling of open chromatin, DNA-binding proteins and nucleosome position. Nat Methods. 2013;10(12):1213–8. doi:10.1038/nmeth.2688.

28. Long T, Bhattacharyya T, Repele A, Naylor M, Nooti S, Krueger S, et al. The contributions of DNA accessibility and transcription factor occupancy to enhancer activity during cellular differentiation. G3 (Bethesda). 2023;doi:10.1093/g3journal/jkad269.

29. Repele A, Krueger S, Bhattacharyya T, Tuineau MY, Manu. The regulatory control of Cebpa enhancers and silencers in the myeloid and red-blood cell lineages. PLOS ONE. 2019;14(6):1–24. doi:10.1371/journal.pone.0217580.

30. Patro R, Duggal G, Love MI, Irizarry RA, Kingsford C. Salmon provides fast and bias-aware quantification of transcript expression. Nature methods. 2017;14(4):417–419. doi:10.1038/nmeth.4197.

31. Anders S, Huber W. Differential expression analysis for sequence count data. Genome Biol. 2010;11(10):R106. doi:10.1186/gb-2010-11-10-r106.

32. Robinson MD, Oshlack A. A scaling normalization method for differential expression analysis of RNA-seq data. Genome Biol. 2010;11(3):R25. doi:10.1186/gb-2010-11-3-r25.

33. Love MI, Huber W, Anders S. Moderated estimation of fold change and dispersion for RNA-seq data with DESeq2. Genome Biology. 2014;15(12):550. doi:10.1186/s13059-014-0550-8.

34. Yu G, Wang LG, Han Y, He QY. clusterProfiler: an R package for comparing biological themes among gene clusters. OMICS. 2012;16(5):284–7. doi:10.1089/omi.2011.0118.

35. Ashburner M, Ball CA, Blake JA, Botstein D, Butler H, Cherry JM, et al. Gene ontology: tool for the unification of biology. The Gene Ontology Consortium. Nat Genet. 2000;25(1):25–29.

